# THE CELLS ARE ALL-RIGHT: Regulation of the *Lefty* genes by separate enhancers in mouse embryonic stem cells

**DOI:** 10.1101/2024.06.14.598870

**Authors:** Tiegh Taylor, Hongyu Vicky Zhu, Sakthi D. Moorthy, Nawrah Khader, Jennifer A. Mitchell

**Affiliations:** Department of Cell and Systems Biology, University of Toronto, Toronto, ON, Canada.; Department of Laboratory Medicine and Pathobiology, University of Toronto, Toronto, ON, Canada.

## Abstract

Enhancers play a critical role in regulating precise gene expression patterns essential for development and cellular identity; however, how gene-enhancer specificity is encoded within the genome is not clearly defined. To investigate how this specificity arises within topologically associated domains (TAD), we performed allele-specific genome editing of sequences surrounding the *Lefty1* and *Lefty2* paralogs in mouse embryonic stem cells. The *Lefty* genes arose from a tandem duplication event and these genes interact with each other in chromosome conformation capture assays which place these genes within the same TAD. Despite their physical proximity, we demonstrate that these genes are primarily regulated by separate enhancer elements. Through CRISPR-Cas9 mediated deletions to remove the intervening chromatin between the *Lefty* genes, we reveal a distance-dependent dosage effect of the *Lefty2* enhancer on *Lefty1* expression. These findings indicate a potential role for both discrete CTCF bound regions of the genome and chromatin distance in insulating gene expression domains in the *Lefty* locus in the absence of architectural insulation.

## INTRODUCTION

The precise regulation of gene expression is a fundamental pillar of developmental biology, guiding the development of complex organisms. Enhancers play a crucial role in orchestrating the finely tuned control of gene expression across diverse tissues and developmental stages. Enhancers function through the coordinated binding by transcription factors which in turn recruit components of regulatory complexes such as mediator; this complex of regulatory factors directly interacts with the RNA polymerase complex recruited to gene promoters to enact a *cis*-regulatory effect (Banerji et al., 1981; Kellis et al., 2014; Nord et al., 2013; Plank & Dean, 2014). Although potentially separated by megabases of DNA, enhancers are brought into close spatial proximity to their gene targets and interact with RNA polymerase at the gene promoter (Lettice, 2003; Tolhuis et al., 2002). These long range interactions can be mediated by the Cohesin complex (Thiecke et al., 2020) which has been proposed to extrude intervening chromatin through its ring-like structure, bringing distal regions into close proximity (Rao et al., 2014). The CCCTC-Binding Factor (CTCF) has been shown to have a stabilizing role on Cohesin and aids in the formation of long-range chromatin interactions (Bell et al., 1999; de Wit et al., 2015), however, the necessity of CTCF in long-range enhancer promoter communication globally has been questioned as the removal of these bound regions can have negligible impact on local chromatin topology and gene expression (Despang et al., 2019; de Wit et al., 2015; Taylor et al., 2022). Global chromatin interactions partition the genome into Topologically Associated Domains (TADs), demarcated by boundaries wherein chromatin interactions occur much more frequently within a TAD than with neighbouring regions (Dixon et al., 2012; Sexton et al., 2012). Regions within TADs show concerted epigenomic signatures and coordinated gene expression highlighting that at a higher level these can act as units of transcriptomic regulation (Le Dily et al., 2014; Nora et al., 2012; Symmons et al., 2016).

During evolution, gene duplication events can give rise to paralogous genes encoding proteins that participate in shared cellular pathways. When both genes are maintained in the genome, the function of each usually diverges, either through modulating gene expression patterns or through changes to the coding sequence that alters protein function (Kuzmin et al., 2022). In tandem duplication events, paralogs remain in close proximity in the genome and are transcribed from the same DNA strand (Lan & Pritchard, 2016). Although these duplicated genes often share expression patterns and appear co-regulated, it has been proposed that after gene duplication there is selection pressure to maintain gene dosage, by reducing paralog expression, so that stochiometric ratios of proteins are maintained (Birchler & Yang, 2022; Gout & Lynch, 2015; Lan & Pritchard, 2016; Qian et al., 2010; Woodhouse et al., 2010). In *Drosophila*, however, paralogs show a high degree of co-regulation, over 75% of paralogs with similar expression patterns display evidence of regulation by shared enhancers or the additive effects of multiple shared enhancers (Levo et al., 2022). Furthermore, enhancer deletion in mouse embryonic stem cells (ESCs) revealed that gene paralogs can be co-regulated by shared enhancers (Moorthy et al., 2017). These data fit with the gene neighborhood theory of gene expression regulation, whereby genes within the same TAD are largely co-regulated by shared enhancers (Ebisuya et al., 2008; Krefting et al., 2018; Loker & Mann, 2022; Symmons et al., 2014), raising the question, how does gene expression control divergence for paralogs in a tandem duplication? Genomic separation of the paralogs appears to be a key factor in expression divergence. Studies have identified a strong correlation in the expression of tandem duplicated genes which are located within one megabase of each other, whereas paralogs which are separated in one species to different chromosomes show much lower correlation in their co-expression patterns (Lan & Pritchard, 2016). This finding holds true when controlling for the age of the duplication as over longer timeframes, duplicated genes are more likely to reside on separate chromosomes (Ghanbarian & Hurst, 2015; Lan & Pritchard, 2016; Sémon & Duret, 2006).

Here, we focus on investigating the regulatory mechanisms controlling *Lefty1* and *Lefty2* transcription. These are paralogous genes central to both stem cell maintenance and the intricate processes of left-right patterning during vertebrate development. In Fugu and Flounder, only a single *Lefty* gene is present which covers the functions of both genes present in other species (Hashimoto et al., 2007). In zebrafish, both paralogs are present but arose from a whole genome duplication event in ray finned fishes (Hashimoto et al., 2007). In mammals, the tandem duplication event leading to the generation of the *Lefty1* and *Lefty2* paralogs was proposed to have occurred separately in mice and humans (Yashiro et al., 2000). Ensembl gene trees predicts that these are separate events (Herrero et al., 2016) and represent an evolutionarily young event (James et al., 2024). Due to the nature of the duplication and gene function, *Lefty2* has been proposed as the ancestral gene (Hashimoto et al., 2007). Given what is known of recently duplicated paralogs, these genes would be expected to have shared regulatory control in mammals and similar expression patterns, however, although both *Lefty* genes are expressed in ESCs and participate in pluripotency maintenance (D.-K. Kim et al., 2014), they diverge in their tissue expression patterns in driving left-right axis patterning during embryonic development, with *Lefty1* primarily being expressed in the prospective floor plate whereas *Lefty2* is expressed in the lateral plate mesoderm (Saijoh et al., 1999).

Despite their joint participation in pluripotency maintenance, and physical presence within the same TAD (Liu et al., 2022), which would suggest enhancer regulatory cross-talk occurs, we determined that transcription of the *Lefty* genes is governed by separate discrete enhancer elements in ESCs. These two enhancers are located upstream of each gene; the *Lefty2* enhancer overlaps the previously identified asymmetric specific enhancer (ASE), whereas the *Lefty1* enhancer is located upstream of the more proximal neural plate enhancer (NPE) and lateral plate mesoderm-specific enhancer (LPE). The ASE, NPE, and LPE were previously identified by injecting mouse pronuclei with enhancer constructs fused to LacZ to identify expression patterns in the developing embryo (Saijoh et al., 1999). Through allele-specific CRISPR-Cas9 mediated deletion, we dissected the elements between the *Lefty* genes that functionally insulate their respective enhancers. By removing 44 kb of intervening chromatin, we show that the *Lefty2* enhancer can regulate *Lefty1* when both are placed in closer proximity. Incremental deletions, which modulate the linear chromatin distance between these two genes, reveal that the *Lefty2* enhancer exerts a distance-dependent dosage effect on *Lefty1* expression, despite the relatively small separation distance of these two genes. These data reveal that although the *Lefty* genes are co-expressed in ESCs and located within the same TAD, they are controlled by separate *cis*-regulatory elements.

## RESULTS

To identify which regions of the *Lefty* locus are responsible for the expression of *Lefty1* and *Lefty2,* we performed CRISPR-Cas9 mediated deletions, using an F1 mouse ESC line (*Mus musculus*^129^ × *Mus castaneus*) to allow for allele-specific deletion, and RNA quantification as described in (Moorthy & Mitchell, 2016). This approach can identify *cis*-regulatory elements by creating heterozygous deletions, avoiding the potentially confounding *trans*-effects of a homozygous deletion that abolishes target gene expression (Moorthy et al., 2017; Zhou et al., 2014). Upstream of either *Lefty* gene are regions we identified as potential enhancers of *Lefty* gene transcription (“Lefty1enh”, “Lefty2enh”, Fig. 1A), as these regions contained multiple binding events for core ESC transcription factors, as well as occupancy of the Med1 subunit of the mediator complex (Fig. 1A). Furthermore, GRO-seq signal was observed on both the positive and negative strand, corresponding to bidirectional transcription (Fig. 1B), a feature often found at enhancer regions (T.-K. Kim et al., 2015). The identified transcription factor bound region upstream of *Lefty2* overlaps part of the previously identified left-side specific ASE enhancer (Saijoh et al., 1999). Intriguingly, both mouse enhancers are also conserved in the human genome, at a sequence level and display enhancer features in human ESCs (Singh et al., 2021).

**Fig 1:**
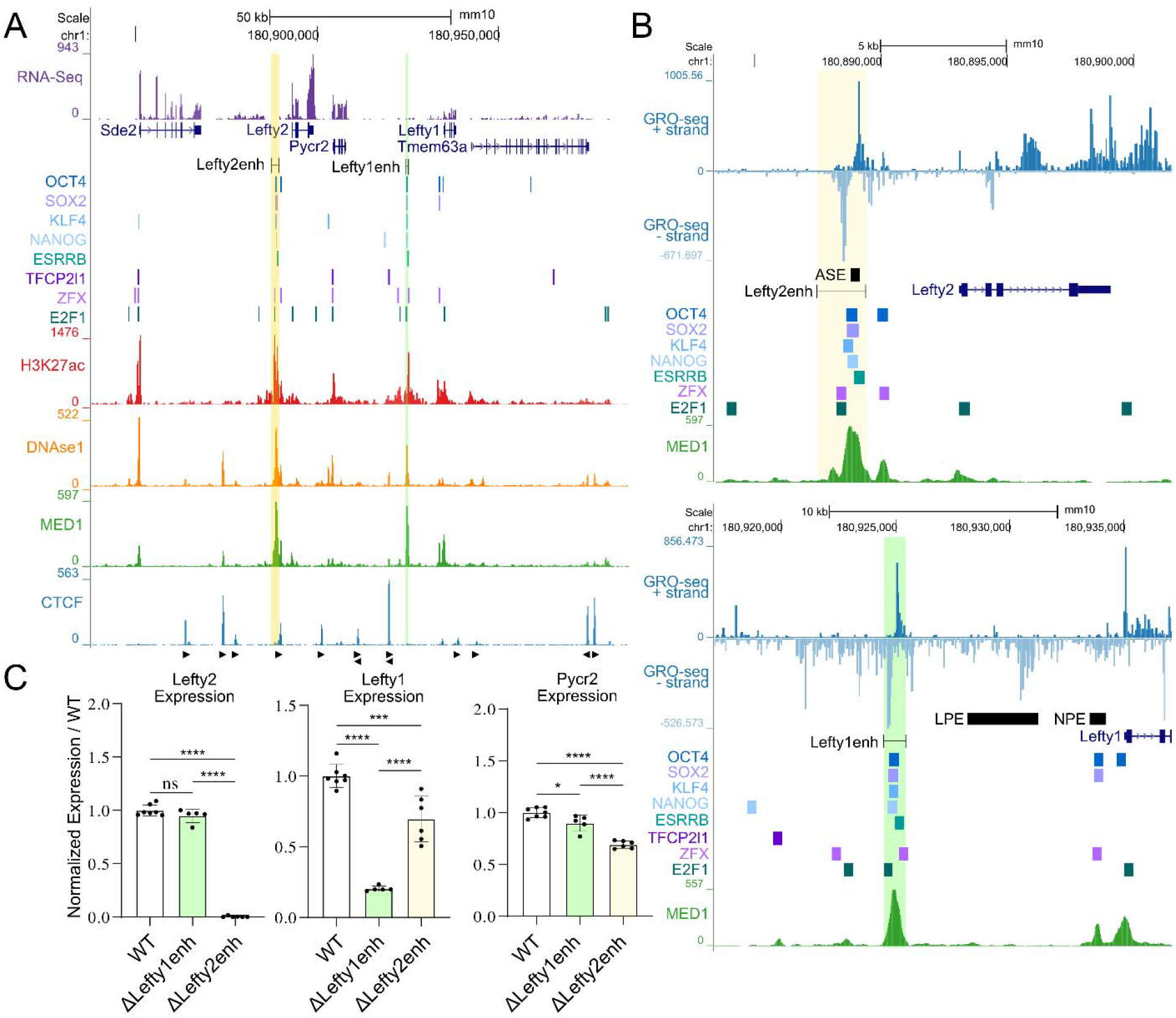
*Lefty1* and *Lefty2* are regulated by separate proximal enhancer regions in mouse embryonic stem cells. A) The *Lefty* locus is displayed on the UCSC genome browser (mm10). The targeted enhancer elements upstream of either *Lefty* gene are highlighted. Transcription factor bound regions derived from mouse ESC ChIP-seq data sets compiled in the CODEX database are shown above ChIP-seq data for H3K27ac, the mediator subunit MED1, and CTCF. Arrows denote CTCF motif orientation. DNAse1-seq data shows accessible chromatin. RNA-seq above the genes shows gene expression in wild-type mouse ESCs. B) Zoomed in view of each *Lefty* enhancer located upstream and proximal to the *Lefty* genes. Strand specific GRO-seq data is shown, bidirectional transcription is a marker of active enhancers. The locations of previously identified enhancers active at embryonic day 8.5 are shown (ASE, LPE, NPE). C) Allele-specific primers detect 129 or *Castaneus* transcripts from *Lefty1*, *Lefty2*, or *Pycr2*. Expression of these genes are normalized to the WT allelic average and represented as a fold change over WT. Deletion of the *Lefty1* enhancer leads to a significant reduction of *Lefty1* transcript abundance with a mild effect on *Pycr2*. Deletion of the *Lefty2* enhancer abolishes *Lefty2* expression while significantly decreasing both *Lefty1* and *Pycr2* transcripts. Error bars represent SD, one way ANOVA significant differences are indicated; (*) P < 0.05, (**) P < 0.01, (***) P < 0.001, (****) P < 0.0001, (ns) not significant.

Removal of the transcription factor bound region upstream of *Lefty2* (ΔLefty2enh) led to a complete abolishment of *Lefty2* transcription while also leading to a significant but minor decrease in transcript abundance for *Lefty1* (∼30%) and *Pycr2* (∼30%) (Fig. 1C), highlighting that this enhancer predominantly regulates *Lefty2* while also contributing in a more minor way to transcription of the other genes in this locus. As *Lefty1* expression was still largely maintained after the loss of the *Lefty2* enhancer, in a separate experiment, a ∼1kb transcription factor bound region ∼9.5kb upstream of *Lefty1* was removed (ΔLefty1enh). Deletion of this region led to a decrease in *Lefty1* expression (∼70%) while having no effect on *Lefty2* and only a minor, yet significant, effect on *Pycr2* (∼10% reduction) (Fig. 1C). Interestingly, we note that the remaining *Lefty1* expression in the *Lefty1* enhancer deleted cell line is similar to the decrease seen when the *Lefty2* enhancer was deleted, highlighting that although the *Lefty2* enhancer may exert some regulatory control over the *Lefty1* gene, it does not act in a functionally redundant or synergistic manner with the *Lefty1* enhancer. In addition, the complete loss of *Lefty2* expression after deletion of the *Lefty2* enhancer revealed that the intact *Lefty1* enhancer is unable to compensate for the loss of the *Lefty2* enhancer (Fig 1C).

We next asked whether the specificity in regulation by the *Lefty* enhancers was due to an architectural insulation of the two genes. Between the *Lefty1* enhancer and *Pycr2* are CTCF bound regions which overlap CTCF motifs both convergent and divergent with *Lefty1*, suggesting this region may act as an architectural boundary (Fig. 2). We evaluated chromatin capture data for this locus to identify if there is evidence of structural insulation between the *Lefty* genes. Hi-TrAC chromatin capture utilizes two linked Tn5 transposases to capture nearby open chromatin and identify long-range chromatin interactions from nearby accessible regions with increased resolution compared to Hi-C (Liu et al., 2022). Both Hi-TrAC and Hi-C data reveals the *Lefty* genes are not architecturally insulated from one another and reside within a shared TAD (Fig. 2, Fig. S1). To evaluate the association between the *Lefty* enhancers and the genes within this TAD, we generated virtual 4C from the Hi-TrAC data with viewpoints centered on either identified enhancer element. The *Lefty2* enhancer shows an association with both the *Lefty1* enhancer and gene, whereas the *Lefty1* enhancer shows an interaction with the *Lefty2* enhancer but minimal signal at the *Lefty2* gene. There is no apparent interaction of the *Lefty2* enhancer with the prominent binding site for CTCF present just upstream of the *Lefty1* enhancer. Noting that the *Lefty* enhancers both physically interact in the nucleus, the question remains as to why these enhancers function separately in the regulation of their cognate *Lefty* genes.

**Fig 2:**
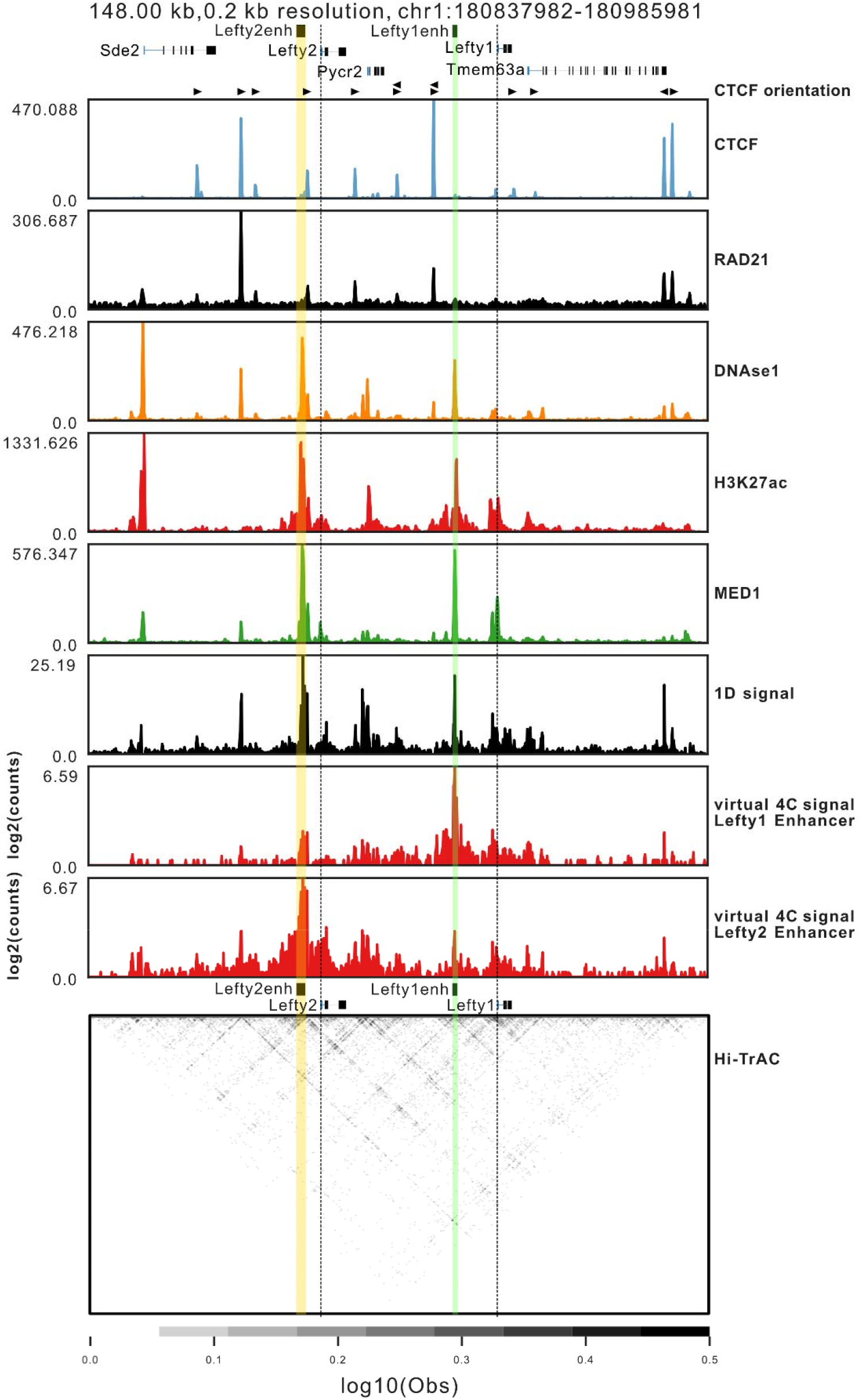
*Lefty* enhancers physically interact in the nucleus and are not architecturally insulated from one another. CLoops2 was used to evaluate Hi-TrAC chromatin capture data, ChIP-seq, and DNase1-seq is presented above the collapsed 1-dimensional chromatin capture signal which highlights regions of frequent interactions. Virtual 4C identifies all interactions with one selected region, in this case tracks for all interactions with the *Lefty1* enhancer or the *Lefty2* enhancer are shown. Dotted lines denote the promoter of both *Lefty* genes while the enhancers have coloured overlays. Both enhancers interact with each other as identified by Hi-TrAC chromatin capture.

Previous studies have suggested enhancer-promoter specificity may act as a layer of cis-regulation in itself, with a ‘lock and key’ type mechanism, preventing off-target enhancer activation, independent of chromatin interactions (Martinez-Ara et al., 2022; Sloutskin et al., 2021). Next, we investigated to what extent the observed specificity of the *Lefty2* enhancer for the *Lefty2* gene promoter may be due to an enhancer-promoter specificity mechanism. To evaluate this possibility, we created a deletion of the intervening chromatin from just upstream of the *Lefty1* promoter region to just downstream of the *Lefty2* enhancer which places the *Lefty1* promoter immediately downstream of the *Lefty2* enhancer (ΔLefty1 to Lefty2enh) (Fig. 3A). This 44kb deletion resulted in an increase in the expression of *Lefty1* (Fig. 3B), significantly above the level of expression observed in WT cells, indicating that there is no apparent enhancer-promoter incompatibility between the *Lefty2* enhancer and the *Lefty1* promoter. In fact, the observed increase in *Lefty1* expression, when under control of the *Lefty2* enhancer, mirrors the observed higher expression levels of *Lefty2* compared to *Lefty1* in ESCs (Fig. 1A). To confirm that there were no other effects on *Lefty1* expression from including a deletion of the sequences between the *Lefty1* enhancer and the more upstream promoter region of *Lefty1*, we created a deletion (ΔLefty1 upstream) which removed this region and confirmed there was no significant difference in expression between this deletion and the previous enhancer deletion (Fig. S2). These findings indicate that the 44kb intervening chromatin region does insulate the *Lefty1* gene from the stronger *Lefty2* enhancer.

**Fig 3:**
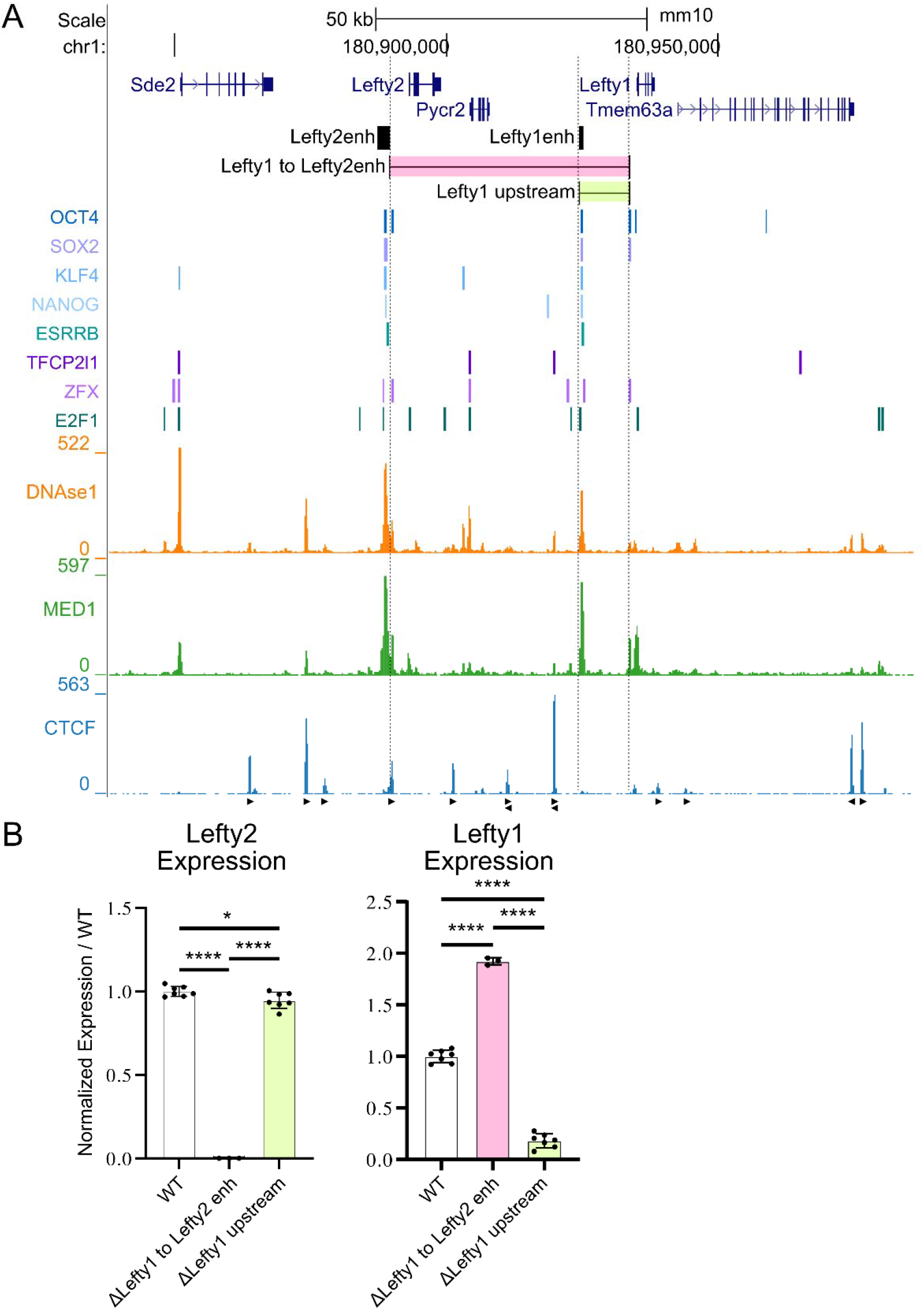
The *Lefty2* enhancer can enhance the expression of *Lefty1* when placed into close linear proximity. A) The *Lefty* locus is displayed on the UCSC genome browser (mm10). The targeted deletions are highlighted along with the previously identified *Lefty* enhancers. Transcription factor bound regions derived from mouse ESC ChIP-seq data sets compiled in the CODEX database are shown above ChIP-seq data for the mediator subunit MED1 and CTCF. Arrows denote CTCF motif orientation. DNAse1-seq data shows accessible chromatin. B) Allele-specific primers detect 129 or *Castaneus* transcripts from *Lefty1* or *Lefty2*. Expression of these genes are normalized to the WT allelic average and represented as a fold change over WT. Deletion of the *Lefty1* enhancer leads to a significant reduction of *Lefty1* transcript. A deletion which places *Lefty1* directly adjacent to the *Lefty2* enhancer causes an increase in *Lefty1* transcript abundance. Error bars represent the SD, one way ANOVA significant differences are indicated. (*) P < 0.05, (**) P < 0.01, (***) P < 0.001, (****) P < 0.0001, (ns) non-significant differences are not marked.

Next we asked, in the absence of architectural insulation or promoter incompatibility, what could cause the transcriptional insulation between the two *Lefty* genes. *Pycr2* is ubiquitously transcribed and needed for proline biosynthesis. We considered the possibility that the *Pycr2* transcription unit is able to act as a sufficient barrier to crosstalk between the two *Lefty* regulatory units. We created deletions of both *Pycr2* alone (ΔPycr2), and in conjunction with deletion of the *Lefty1* upstream region (ΔLefty1 upstream + Pycr2), to see if the absence of the *Pycr2* transcription unit would cause an increase in transcription of *Lefty1* (Fig 4A). However, we observed no effect on the expression of *Lefty1* or *Lefty2* when *Pycr2* is deleted alone, and when deleted in compound with the *Lefty1* enhancer, there was no rescue of *Lefty1* expression compared to deletion of the *Lefty1* enhancer alone (Fig 4B).

**Fig 4:**
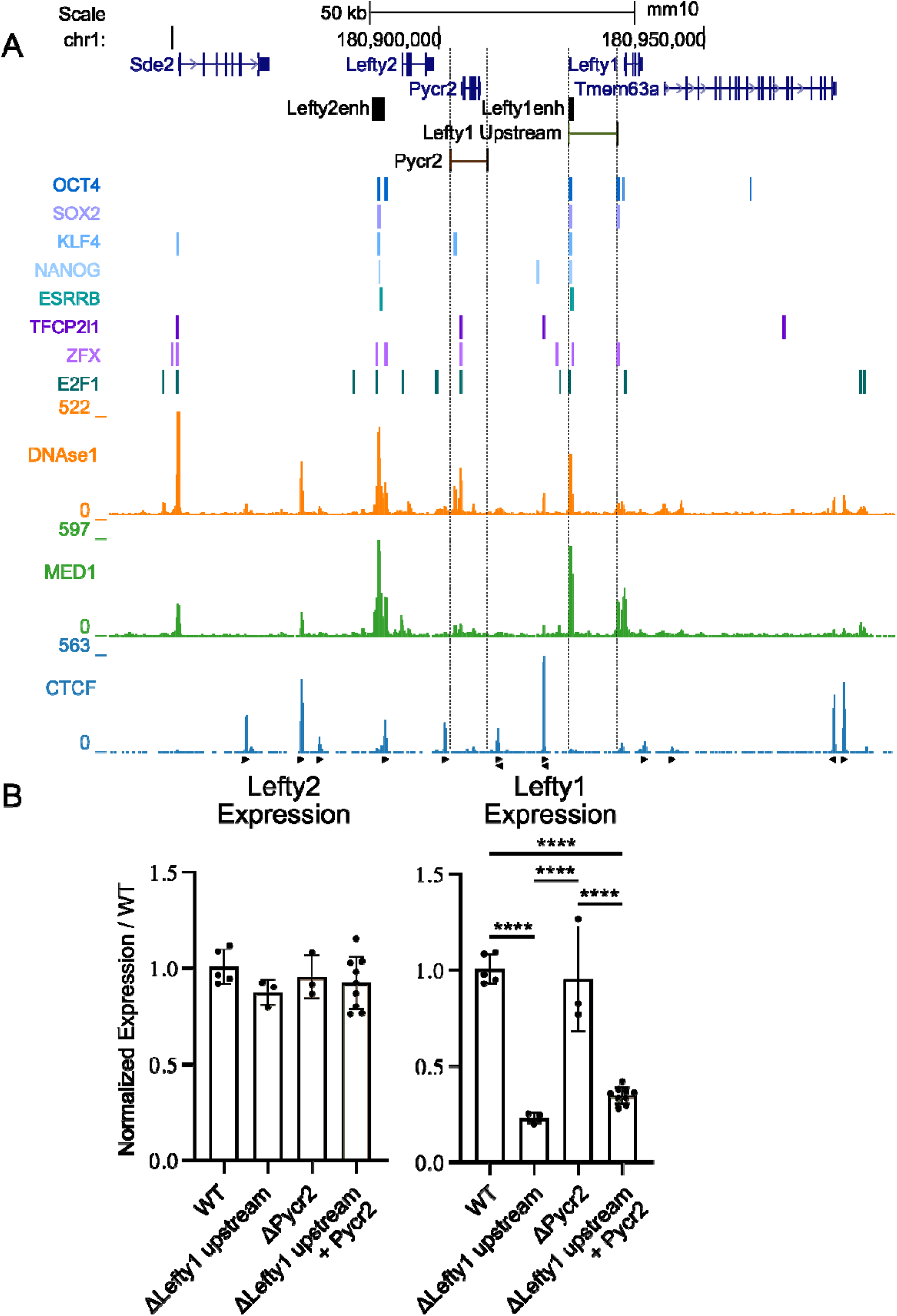
The universally transcribed *Pycr2* gene is not responsible for the transcriptional insulation of the *Lefty* genes. A) The *Lefty* locus is displayed on the UCSC genome browser (mm10). The targeted deletions are highlighted along with the previously identified *Lefty* enhancers. Transcription factor bound regions derived from mouse ESC ChIP-seq data sets compiled in the CODEX database are shown above ChIP-seq data for the mediator subunit MED1 and CTCF. Arrows denote CTCF motif orientation. DNAse1-seq data shows accessible chromatin. B) Allele-specific primers detect 129 or *Castaneus* transcripts from *Lefty1*, *Lefty2*. Expression of these genes are normalized to the WT allelic average and represented as a fold change over WT. Deletion of *Pycr2* alone or in conjunction with the region upstream of *Lefty1* overlapping the enhancer do not significantly affect *Lefty1* expression as compared to WT or parent enhancer deleted clone. Error bars represent the SD, one way ANOVA significant differences are indicated. (*) P < 0.05, (**) P < 0.01, (***) P < 0.001, (****) P < 0.0001, non-significant differences are not marked.

To identify what element or elements are responsible for the insulation of *Lefty1* from the *Lefty2* enhancer, we performed further deletion experiments to remove portions of the intervening chromatin between these two genes. We performed a stepwise deletion approach, removing larger and larger portions of the chromatin that separates the genes. We created three deletions, 1) removing ∼31kb, placing *Lefty1* downstream of *Lefty2* (ΔL1-31kb), 2) removing ∼24.5kb placing *Lefty1* downstream of *Pycr2* (ΔL1-24.5kb), and finally 3) removal of ∼14.5kb and the most prominent CTCF bound region upstream of the *Lefty1* enhancer (ΔL1-14.5kb); these are each compared with the previous *Lefty1* upstream deletion which removed ∼9kb and the *Lefty1* enhancer (ΔLefty1 upstream) (Fig 5A). We observed an increase in the expression of *Lefty1* as larger portions of the intervening chromatin were removed. In the ΔL1-14.5kb deletion, we observe a great deal of variation between the different independent clonal isolates such that, although the average decrease in expression was 55% for these clones, these levels were not significantly different from either the enhancer deleted ΔLefty1 upstream clones or the wild-type expression levels (Fig 5B). The clonal variation did not appear to correspond to any differences in the deletion boundaries inherent to CRISPR-Cas9 based deletions (Fig. S3, Table S1). As larger deletions were made, shortening the separation distance between *Lefty1* and the *Lefty2* enhancer, we observed further increased *Lefty1* expression which reached wild-type levels after the 31kb deletion (ΔL1-31kb deletion). In all cases, there was no significant change in the expression of *Lefty2* across the clones, highlighting that the *Lefty2* enhancer crosstalk regulating *Lefty1* does not affect its efficacy in enhancing the expression of *Lefty2*.

**Fig 5:**
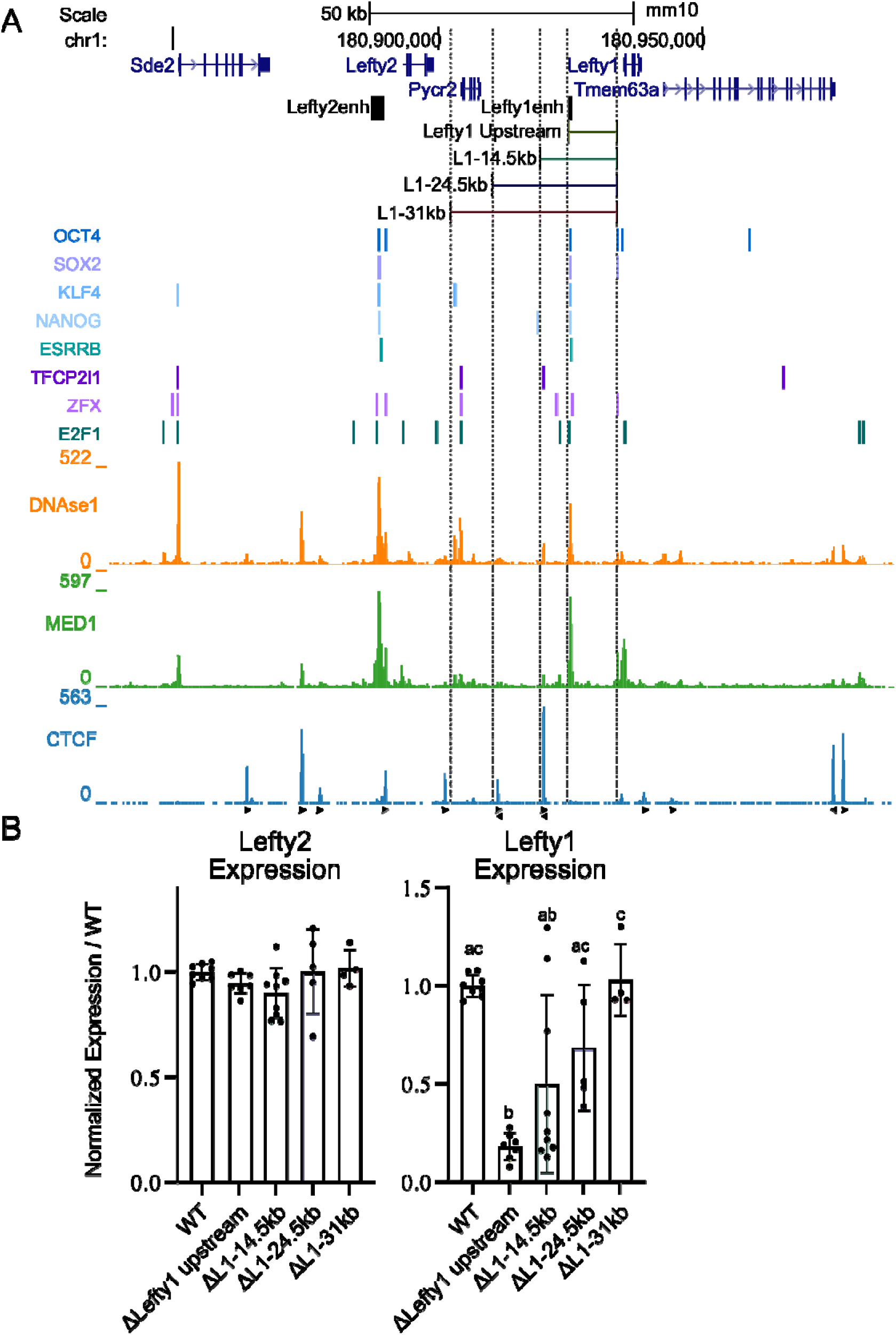
Regulatory capacity of the *Lefty2* enhancer on *Lefty1* expression is modified by the intervening chromatin. A) The *Lefty* locus is displayed on the UCSC genome browser (mm10). The targeted deletions are highlighted along with the previously identified *Lefty* enhancers. Transcription factor bound regions derived from mouse ESC ChIP-seq data sets compiled in the CODEX database are shown above ChIP-seq data for the mediator subunit MED1 and CTCF. Arrows denote CTCF motif orientation. DNAse1-seq data shows accessible chromatin. B) Allele-specific primers detect 129 or *Castaneus* transcripts from *Lefty1*, *Lefty2*. Expression of these genes are normalized to the WT allelic average and represented as a fold change over WT. A dosage dependent effect of *Lefty2* enhancer crosstalk on *Lefty1* transcription is observed as more intervening chromatin is removed. Error bars represent the SD, one way ANOVA significant groups with P < 0.05 are indicated with letters.

## DISCUSSION

Using CRISPR-Cas9 mediated genome editing to remove sections of the *Lefty* locus, we identified the *cis*-regulatory elements responsible for *Lefty1* and *Lefty2* transcriptional control in ESCs, as well as identifying a curious case of transcriptional insulation in the absence of obvious architectural insulation. We determined that the *Lefty1* and *Lefty2* genes are both primarily regulated by separate enhancer elements located just upstream of either gene. The entirety of *Lefty2* transcriptional regulation in ESCs can be ascribed to a ∼1.9kb region located ∼3.7kb upstream, that overlaps the ASE. Conversely, although the majority of *Lefty1* transcriptional regulation is due to a ∼1kb transcription factor bound region located ∼9.5kb upstream, there is an additive effect on *Lefty1* transcription from the *Lefty2* enhancer which accounts for ∼30% of *Lefty1* expression. The region we identify as the required *cis*-regulatory element for *Lefty1* differs from the previously identified more proximal ESC enhancer element (located 1.3kb upstream of *Lefty1*); identified by luciferase reporter assays rather than genetic perturbation (Nakatake et al., 2006). The ESC *Lefty2* enhancer overlaps the previously established ASE responsible for driving *Lefty2* expression in the lateral plate mesoderm during embryonic development (Saijoh et al., 1999), indicating this element is active in more than one context. In addition, we show that these two genes and their cognate ESC enhancers interact in conformation capture assays that show no evidence of architectural insulation between the two genes. Despite this apparent lack of architectural insulation, we find that the 44kb of chromatin separating these two genes does transcriptionally insulate the *Lefty2* enhancer from the *Lefty1* gene. Furthermore, partial deletions of this region tuned *Lefty1* transcript levels in relation to the separation distance from the *Lefty2* enhancer, suggesting insulation at this locus is highly distributed, similar to what has been observed for some TAD boundaries (Huang et al., 2021). Other studies have determined that linear chromatin distance does impact enhancer activity when moving the β-globin enhancer (LCR) further away from its promoter when inserted at ectopic loci (Rinzema et al., 2022). Distance also plays a more pronounced role in regulating the activity of weaker enhancers, as a reporter assay testing the effect of distance on enhancer activity found that the activity of the *Nanog* enhancer rapidly dropped off as it was shifted from 25kb away from the target gene to 75kb (Henry Thomas et al., 2023). In *Drosophila*, shifting the GMR enhancer from a few hundred base pairs away to 3kb away completely attenuated its ability to activate a target *hsp70* promoter (Bateman & Johnson, 2022). Our study reveals the distance dependent insulation of the *Lefty1* gene from the stronger *Lefty2* enhancer which may have been involved in the functional divergence of these gene paralogs by allowing for downregulated *Lefty1* dosage after the duplication event.

Insulation at a number of other loci has been linked to CTCF binding events (Bell & Felsenfeld, 2000; Flavahan et al., 2016; Guo et al., 2015) and convergent CTCF motifs are enriched at adjacent TAD boundaries, and required for TAD boundary maintenance (Dixon et al., 2012; Nora et al., 2017; Williamson et al., 2019; Zuin et al., 2014). Given this role for CTCF at other regions of the genome, we were surprised to find that deletion of the most prominent CTCF bound region that separates the two *Lefty* genes did not abolish insulation in all clones but instead caused clonal variation in *Lefty1* expression levels, ranging from completely insulated to a restoration of wild-type expression levels, whereas *Lefty2* levels were unaffected. This finding indicates that the prominent CTCF bound region between *Lefty1* and *Pycr2* confers robustness to the insulation between the *Lefty* paralogs but this insulation can be maintained in its absence. Two additional regions between the *Lefty* genes display CTCF binding and removal of one of these regions with the 24.5kb deletion did relieve additional insulation of the Lefty1 gene, while still displaying clonal variation. Interestingly, the largest deletion of 31kb that leaves both genes intact did not remove an additional CTCF bound region but did abolish insulation, bringing the expression of *Lefty1* to wild-type levels. Together these findings suggest a role for both discrete CTCF bound regions of the genome and chromatin distance in insulating gene expression domains in the *Lefty* locus. Genome-wide insulator boundaries do not seem to exist as discrete elements; a striking ∼20% of TAD boundaries remain stable upon the degradation of CTCF (Nora et al., 2017) and investigations into the effects of removing TAD boundaries have mapped insulator regions of up to 80kb in size which are responsible for boundary maintenance (Rajderkar et al., 2023). Other studies have shown that modulating the distance between enhancer-promoter pairs in endogenous and artificial loci does modulate enhancer activity, indicating chromatin distance can have an insulator effect (Bateman & Johnson, 2022; Rinzema et al., 2022).

Highlighting the complex interplay of chromatin architecture and gene regulation, studies which have temporally degraded architectural factors, such as CTCF or the RAD21 subunit of the Cohesin complex, identified surprisingly few changes in global transcription even upon a near complete loss of TAD structures (Hsieh et al., 2022; Nora et al., 2017; Rao et al., 2017). However, it is difficult to draw concrete conclusions from these data about how CTCF degradation impacts *Lefty* expression, as we see conflicting results from different studies. For example, after 24-48 hours of CTCF ablation using the Auxin-inducible degron system, RNA-seq data shows a significant increase in expression of both *Lefty1* and *Lefty2* (Nora et al., 2017), which could be due to a loss of insulation. Another study degrading CTCF over 3, 12, or 24hrs found a modest but significant increase in *Lefty1* at 24hrs by RNA-seq, however, the same study measured a steady and significant decrease in *Lefty1* nascent transcript by mNET-seq at 3hr, which grew more severe after 24hr of CTCF degradation (Hsieh et al., 2022). These confounding results highlight issues with global protein degradation as the *trans*-regulatory environment is profoundly affected, causing both direct and indirect gene expression changes.

Additional complex mechanisms can tune gene expression levels, for example promoters can compete for enhancer activity which may act to dilute the activity of an enhancer away from other nearby genes (Bateman & Johnson, 2022; Oh et al., 2021), although in the α-globin locus one enhancer acts to regulate multiple genes by forming a single complex hub, arguing competition may not occur in all cases (Oudelaar et al., 2019). We saw evidence for promoter competition at the *Lefty* locus as the *Lefty2* enhancer exerts the greatest effect on *Lefty1* expression when the *Lefty2* gene was also deleted in the 44kb deletion. In this deletion, *Lefty1* expression levels were observed to increase above the WT levels for this gene due to control by the stronger *Lefty2* enhancer. It is also possible, however, that this apparent promoter competition is simply the result of positioning the *Lefty1* gene even closer to the *Lefty2* enhancer as we are comparing a 31kb and 44kb deletion in this case.

We conclude that although the *Lefty* enhancers contact each other as well as both *Lefty* genes, as detected by chromatin conformation capture methods, there is a distance and CTCF regulated insulation between the two genes that allows for reduced *Lefty1* expression. Other studies have shown that tethering an enhancer to an off-target gene can lead to aberrant transcription, opposing our findings (Bartman et al., 2016; Peslak et al., 2023), however, there may be differences in the magnitudes of the chromatin interaction in these two cases. It has also been observed that by shifting an enhancer and it’s cognate gene closer together, one can increase and maintain gene expression over time in an artificial locus (Rinzema et al., 2022). In addition, distance may have enhancer specific effects on the cooperativity between elements (Henry Thomas et al., 2023). Similar to the cooperation observed between multiple CTCF sites that maintain architecture in the *Shh* locus (Williamson et al., 2019), or the ability for a stronger enhancer to be less affected by CTCF insulation (Chakraborty et al., 2023), we observed that CTCF bound regions between the *Lefty* genes played a partial but incomplete role in insulating the expression of these paralogs.

## MATERIALS AND METHODS

### Cell culture

Mouse F1 ESCs (M. musculus129 × M. castaneus; female cells obtained from Barbara Panning) were cultured on 0.1% gelatin-coated plates in ES medium (DMEM containing 15% FBS, 0.1 mM MEM nonessential amino acids, 1 mM sodium pyruvate, 2 mM GlutaMAX, 0.1 mM 2-mercaptoethanol, 1000 U/mL LIF, 3 µM CHIR99021 [GSK3β inhibitor; Biovision], 1 µM PD0325901 [MEK inhibitor; Invitrogen]), which maintains mouse ESCs in a pluripotent state in the absence of a feeder layer (Mlynarczyk-Evans et al., 2006; Ying et al., 2008).

### Cas9-mediated deletion

Cas9-mediated deletions were carried out as previously described (Moorthy & Mitchell, 2016; Zhou et al., 2014). Cas9 targeting guide RNAs (gRNAs) were selected flanking the desired region identified for deletion (Supplemental Table S2). For select cases of allele-specific targeting, gRNA pairs were designed so that at least one gRNA overlapped a SNP to specifically target the M. musculus^129^ allele. On- and off-target specificities of the gRNAs were calculated as described in (Doench et al., 2016; Hsu et al., 2013) to choose optimal guides with a minimal score of 50 for off-target effects. On-target specificity is confirmed by multiple sequence alignments confirming the lack of a functional PAM and/or multiple mismatches in the 3’ end of the Guide RNA. Guide RNA plasmids were assembled with gRNA sequences using the protocol described by (Mali et al., 2013) into (Addgene 41824)or ordered as custom synthesized plasmids containing the entire gRNA expression cassette in a minimal backbone (IDT).

F1 mouse ESCs were transfected with 5 µg of each of the 5′ gRNA(s), 3′ gRNA(s), and pCas9_GFP (Addgene 44719) (Ding et al., 2013) plasmids using the neon or neon nxt transfection systems (Life Technologies). Forty-eight hours after transfection, GFP-positive cells were isolated on a BD FACSAria. Ten-thousand GFP-positive cells were seeded on 10-cm gelatinized culture plates and grown for 5–6 d until large individual colonies formed. Individual colonies were picked and propagated for genotyping and gene expression analysis as previously described (Moorthy & Mitchell, 2016; Zhou et al., 2014). Genotyping of the deletions was performed by amplifying products internal to and surrounding the target deletion. All deleted clones identified from the initial screen were sequenced across the deletion; SNPs confirmed allele specificity of the deletion (Supplemental Table S1).

### RNA isolation and gene expression analysis by RT-qPCR

Total RNA was purified from single wells of >85% confluent six-well plates using the RNeasy Plus mini kit (Qiagen), and an additional DNase I step was used to remove genomic DNA. RNA was reverse-transcribed with random primers using the high-capacity cDNA synthesis kit (Thermo Fisher Scientific). Gene expression was detected by allele-specific primers that specifically amplified either the musculus or castaneus allele as described in (Moorthy & Mitchell, 2016). The standard curve method was used to calculate expression levels using F1 mouse ESC genomic DNA to generate the standard curves. Levels of Sdha RNA were used to normalize expression values. Primer sequences are listed in Supplemental Table S3.

### Hi-TrAC Reanalysis

Processed Hi-TrAC .bedpe files for E14 mouse ESCs were obtained from GSE180175 (Liu et al., 2022). cLoops2 was used to create profiles and heatmaps in Figure 2 (Cao et al., 2022) (https://github.com/YaqiangCao/cLoops2).

### Hi-C Reanalysis

Published Hi-C data (Bonev et al., 2017) were downloaded from GEO (GSM2533818–2533821 for mouse ESC DpnII and reanalyzed using the FAN-C toolbox (Kruse et al., 2020), entailing read-mapping, filtering out technical artifacts, mapping to restriction fragment space, binning, matrix normalization, ratio-based comparison, and visualization.

### UCSC data visualization

Mouse ESC ChIP-seq, DNAse-seq, RNA-seq data sets, and associated peak files were obtained from the CODEX database (Sánchez-Castillo et al., 2015). GRO-seq datasets were obtained from GSE186687 (Tafessu et al., 2023)

## ACKNOWLEDGEMENTS

We thank Barbara Panning for sharing resources and members of the Mitchell lab for their guidance and feedback on this project. Work in the Mitchell laboratory was supported by the Canadian Institutes of Health Research (FRN 153186), the National Institutes of Health (R01-HG010045-01), the Canada Foundation for Innovation, and the Ontario Ministry of Research and Innovation. This research was enabled in part by support provided by Compute Ontario (https://www.computeontario.ca/) and the Digital Research Alliance of Canada (https://www.alliancecan.ca). H.V.Z., is supported by the Natural Sciences and Engineering and Research Council (NSERC) of Canada’s Undergraduate Student Research Award (USRA).

## Author Contributions

T.T., S.D.M., and J.A.M. conceived and designed the study. T.T., H.V.Z., N.K., and S.D.M. created ESC deletion lines and analyzed gene expression. T.T. performed analysis of Hi-TrAC and Hi-C datasets. All authors contributed to writing the manuscript.

## Notes

### Competing Interest Statement

The authors have declared no competing interest.

