## Supplemental data for "THE CELLS ARE ALL-RIGHT: Regulation of the *Lefty* genes by separate enhancers in mouse embryonic stem cells"

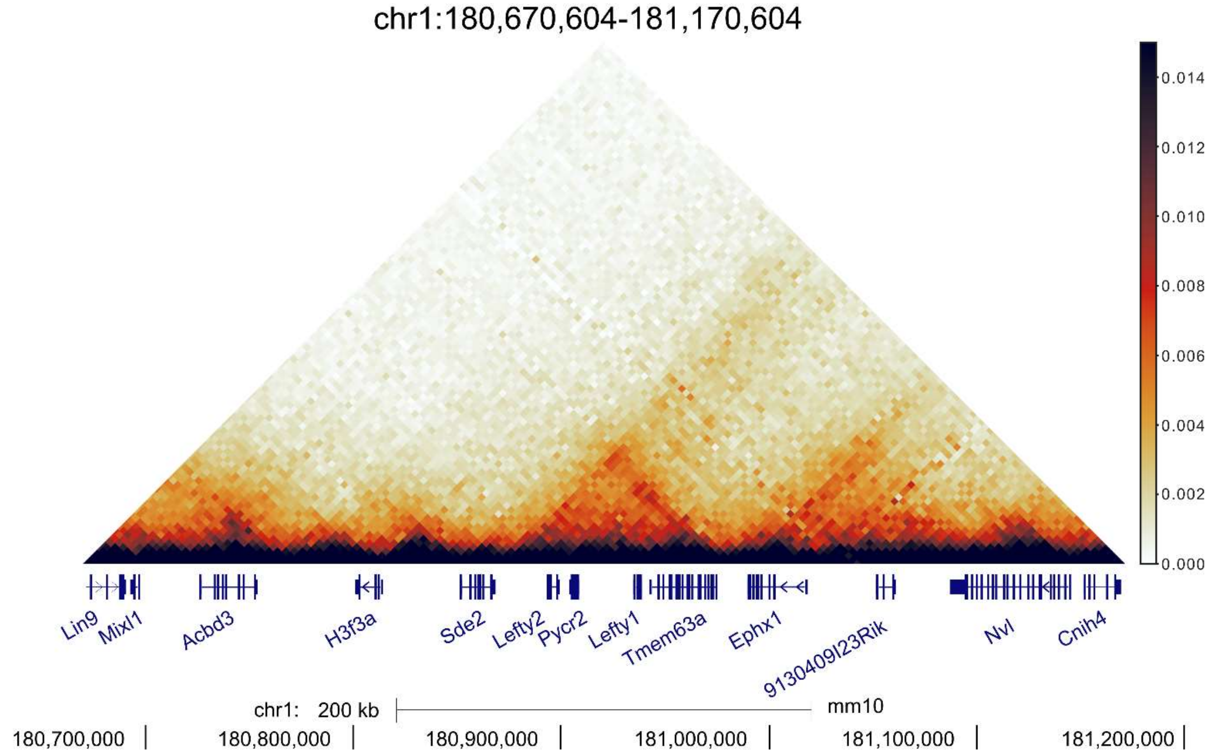


**Supplemental Figure S1:** Hi-C data from mouse ESCs (acquired from (Bonev et al., 2017)) indicating the frequency of occurring interactions surrounding the *Lefty* locus at a 4kb resolution from chr1:180,670,604-181,170,604 mm10. Genes are represented along the bottom of the heatmap. The *Lefty* genes reside within a shared interacting region.

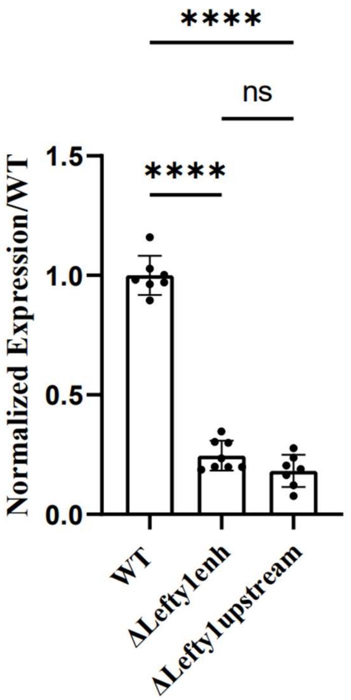


**Supplemental Figure S2:** Comparison of *Lefty1* expression in Lefty enhancer deleted cells vs Lefty1 upstream deleted cells. No Significant difference in expression is observed in *Lefty1* expression between the two deletions, both are significantly reduced in expression as compared to wild-type cells. Error bars represent the SD, one way ANOVA significant differences are indicated. (*) $P < 0.05$, (**) $P < 0.01$, (***) $P < 0.001$, (****) $P < 0.0001$, (ns) not significant.

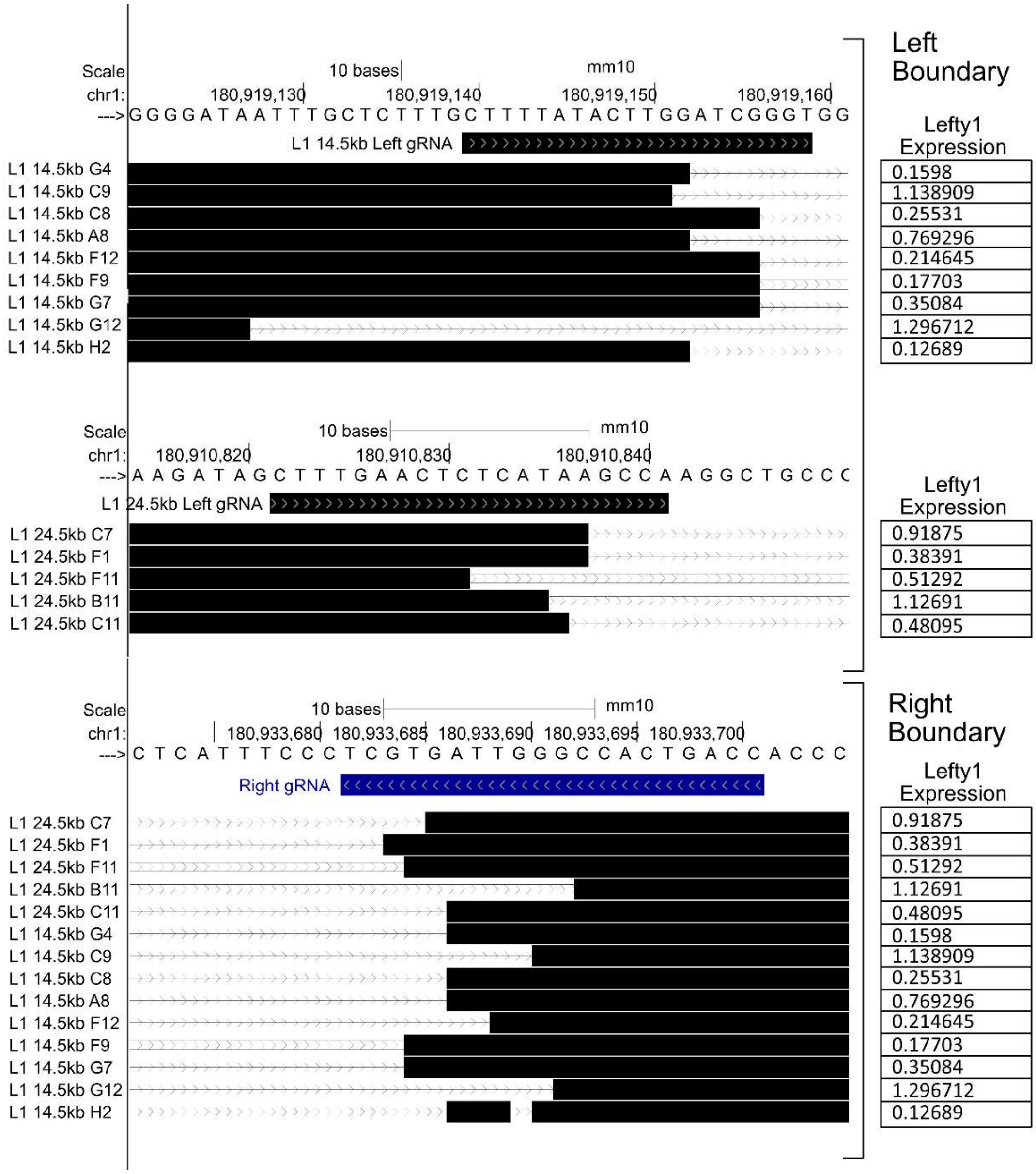


**Supplemental Figure S3:** UCSC browser view of the deletion boundaries which showed highly variable *Lefty1* expression. The precise breakpoints of L1 14.5kb and L1 24.5kb deletions varied by <32bp, differences in deletions boundaries do not seem to correlate with the clonal variability in *Lefty* expression which was observed. Values of *Lefty1* expression by clone are shown in the table on the right, the values are shown as a proportion of wild-type expression. Clone names are shown beside the left or right boundary of the sequenced deletion, dark bars show the remaining aligned genomic sequence whereas the thin lines represent the removed region.

### S1 Table: Sequences Across CRISPR-Cas9 Mediated Deletions

Clone name denotes deleted allele in 129/Cast Cells

| Deletion | Clone | Sequence |
|---|---|---|
| ΔLefty1enh | D5/+ | TTAGGACCCTGCTTTGAACCTNCTGGGTTCCAGGAGAAGCTGATGC<br>CCTCACACAGATATACATGCAGGCAAACCATCAACGCACACAAGAG<br>AAAAAAATAAAGGTAAGATCTTACCACGTAGCTCTGGCCGGCTCGG<br>AAGTTCCTTGTAAATCCGACCGGCTTGAAACTCACAGAAATTCATCT<br>GCCTTTGCCTCCCAAGTCCTGGGTTAAAACTTCTGCACTACTATGCT<br>CATTTTATTTTGAATTTTGATACAAGGTCTCACTGTAGCCAGCTAAG<br>GCTCGcCTCCAGTTCACTATGTAGCTAAAGATGGCCTTGGACTTCTG<br>GTTCTCTTGCCTCTGCCTTCCAAGTGCTGGAATCGCAAGCCCaTATC<br>GCCATTCCCTGCTTCTATAATTAACTTTACAGACTTtACTTTTGGAgG<br>GTTTCGACTAATGCAGCTCAcTtTATAATAGAAGTTTTtCAAGGATTA<br>TCACTATTTTGGAGGTCGCCCAGCAAGGGATGGAGGGTCTCAAACA<br>GAAAATGAGACCCAGTGAGACCATAC\|GGCCCTGTGAGCTCCTGACC<br>CTAGCCCAGAGCACGGTACAAGCCATGTGAATCCTGAGTTTAGTTTT<br>AGATGTGTGTGTCAGGAGNNCCCTTTTGGGTAAA |
| | D6/+ | TTAGNCCCTGCTTTGTATACTTCTNCGTTCCAGGTAGTANCTGATGC<br>CCTCACACAGATATACATGCAGGCAAACCATCAAAGCACATAAGAT<br>AAAAAAATAAAGGTAAGATCTTACCATGTAGCTCTGGCTTGCTTGG<br>AAGTTCCTTGTAAATCCGACTGGCTTGAAACTCACAGAAATTCATCT<br>GCCTTTGCCTCCCAAGTCCTGGGTTAAAACTTCTGCACTACTATGCT<br>CATTTTATTTTGAATTTTGATACAAGGTCTCACTGTAGCCAGCTAAG<br>GCTCGcCTCCAGTTCACTATGTAGCTAAAGATGGCCTTGGACTTCTG<br>GTTCTCTTGCCTCTGCCTTCCAAGTGCTGGAATCGCAAGCCCaTATC<br>GCCATTCCCTGCTTCTATAATTAACTTTACAGACTTtACTTTTGGAgG<br>GTTTCGACTAATGCAGCTCAcTtTATAATAGAAGTTTTtCAAGGATTA<br>TCACTATTTTGGAGGTCGCCCAGCAAGGGATGGAGGGTCTCAAACA<br>GAAAATGAGACCCAGTGAG\|CTCCTGACCCTAGCCCAGAGCACGGT<br>ACAAGCCATGTGAATCCTGAGTTTAGTTTTAGATGTGTGTGTCAGGA<br>GACCCCTTTGGGAAAA |
| ΔLefty2enh | 22B/+ | CATGGGCTCTGGGGTGACCTCTAGGGGCGTTAAATTGGGAAGCATA<br>GGTGATCCACACTGAGAAACTAAGCCCCCTTCCCAATTTAGGGA\|GG<br>TTCAAAACCTCCCATAAATCCAACTCCAGGGAATTCAACAGCCTCT<br>CCTGGCCGTCCTGGGCACCCGCgCACACGTGGTAGACAGCCACGCA<br>GATACGCTTAAATTAAaAGTAAATTTAAAAGAATACAGAAATCACA<br>TtGGAAATTTTCCTGGGATGTAACTCAGCTGGCCTAGCTCCAGATCC<br>TGGGTTCAGATGAGCGCTGCAGACAAAGAAGAGTGGTGCTATGTCT<br>GCCTCCTGTTCTACCTACCTCCTCCTCCCTTTATGGGAGGTTCAAAA<br>CCTCCCATAAATCCAACTCCAGGGAATTCAACAGCCTCTCCTGGCC<br>GTCCTGGGCACCCGCGCACACGTGGTAGACAGCCACGCAGATACGC<br>TTAAATTAAAAGTAAATTTAAAAGAATACAGAAATCACATTGGAAA<br>TTTTCCTGGGATGTAACTCAGCTGGCCTAGCTCCAGATCCTGGGTTC<br>AGATGCAGCGCTGCAGACAAAGAAGAAAGAAGAAAGCATTAGCC<br>CTAGAAT |
| | +/33C | TACGAAATGAGAGATGGAGGCATTCTAGGGCTAATGCTTTCTTCTTT<br>CTTCTTTGTCTGCAGCGCTGCATCTGAACCCAGGATCTGGAGCTAGG<br>CCAGCTGAGTTACATCCCAGGAAAATTTCCgATGTGATTTCTGTATT<br>CTTTTAAATTTACTTTTAATTTAAGtGTATCTGCGTGGCTGTCTACCA<br>CGTGTGtGCGGGTGCCCAGGACGGCCAGGAGAGGCTGTTGAATTCC<br>CTGGAGTTGGATTTATGGGAGGTTTTGAACCACCTAATTGGGAAGG<br>GGGCTTAGTTTCTCAGTGTGGATCACCTATGCACGCCCCTAGAGGTC<br>ACCCCAGGAGTGCTCTGGGGAAGGGTTCtCCAGTTCAGCCTCAGGGA<br>TACGTGCATAGGTGATCCACACTGATATACTTTCCCTTCCCATG\|AGG |

<table>
<tr><td></td><td></td><td>TGGTTCGAACCTCCCAAAATTTTACTCCGGGAATTCAACAGCCTCTC<br>CTGGGCGTCCTGGGCCCCGCaCACCCGTGGTATACGCCACGCACATC<br>CCCTTAAATTAAaAGTAAATTTAAAAGAATACAGAAATCCATcGGAA<br>ATTTTCTGGGATGTAACTACTGCCTAGCTCCGATCGGGTTCGATGCG<br>CGCTGCACACCAGAAGAAAGAAGAAACCTTAGCCCTAGAATGCCTC<br>CTCTTCTTTCAATCCTGTCCCTCCCACTCTCCCCCCAAAA</td></tr>
<tr><td></td><td>54B/+</td><td>TACAGAAAATAGAGAGATGGAGGCATTCTAGGGCTAATGCTTTCTT<br>CTTTCTTCTTTGTCTGCAGCGCTGCATCTGAACCCAGGATCTGGAGC<br>TAGGCCAGCTGAGTTACATCCCAGGAAAATTTCCaATGTGATTTCTG<br>TATTCTTTTAAATTTACTTTTAATTTAAGcGTATCTGCGTGGCTGTCT<br>ACCACGTGTGcGCGGGTGCCCAGGACGGCCAGGAGAGGCTGTTGAA<br>TTCCCTGGAGTTGGATTTATGGGAGGTTTTGAACC|<u>TCCCTAAATTGG</u><br><u>G</u>AAGGGGGCTTAGTTTCTCAGTGTGGATCACCTATGCACGCCCCTA<br>GAGGTCACCCCAGGAGTGCTCTGGGGAAGGGTTCgCCAATTCAGCC<br>TCAGGGATAATTGCACAGCTTCTCCTCTCCGTCCTGCTCATCCCTG<br>CACAACTGGGGGAGGACCATTTCTACCCCCTTAACTCCCCCGGGAA<br>TTTAGAGAGTCTACAGGTCTTCCTGGGCAATTTTTTCCTGGGATAAA<br>ACTCCGCTAAGCCCCTCTACTGATCGTGAAATTCAGATGCAAACAG<br>AAATCAATTGGAAAATTTTCTGGGGCGTTACTCCCTGGCTGCCTCCG<br>ACTTGGGTCTCGACGCTGTCCCCACAACAAGAATAAAGAACACAGG<br>CTTGCCCTAGGAATGCCCCCTCTTCCTCTAAAATCCCGTTCCCCACC<br>TCCCCCCCCCCCACAAATT</td></tr>
<tr><td></td><td>+/87C</td><td>CCGGAATAGAGAGATGGAGGCATTCTAGGGCTAATGCTTTCTTCTTT<br>CTTCTTTGTCTGCAGCGCTGCATCTGAACCCAGGATCTGGAGCTAGG<br>CCAGCTGAGTTACATCCCAGGAAAATTTCCgATGTGATTTCTGTATT<br>CTTTTAAATTTACTTTTAATTTAAGtGTATCTGCGTGGCTGTCTACCA<br>CGTGTGtGCGGGTGCCCAGGACGGCCAGGAGAGGCTGTTGAATTCC<br>CTGGAGTTGGATTTATGGGAGGT|CCTATGCACGCCCCTAGAGGTCA<br>CCCCAGGAGTGCTCTGGGGAAGGGTTCtCCAGTTCAGCCTCAGGGAA<br>ATTGAACAGCCTCTCCCGGCCGTCCTGGGCACCCGCaCACAACGTGG<br>TACAGGGCCACGCAGATCCTCTTAAATTAAGAGTAAATTTAAAAGA<br>ATACAGAAATCACATcGGATATTTTCCTGGGATGTAACTCACCTGGC<br>CTAGCTCCGGATCCGGGGTTCAGATGCAGCGCTGCAGACGGAGAAG<br>AAAGAAGAACGCATTAGCCCTAGAATTGCCTCCGTCTTCTTCTAAAT<br>CCCTGTCCCCTCAACTACCTCCTCCTCCCA</td></tr>
<tr><td></td><td>+/92C</td><td>TGTGGCTCTGGGGTGACCTCTAGGGGCGTGCATAGGTGATCCACAC<br>TGAGAAACTAAGCCCCCTTCCCAATT|AGGTGGTTCAAAACCTCCCA<br>TAAATCCAACTCCAGGGAATTCAACAGCCTCTCCTGGCCGTCCTGG<br>GCACCCGCaCACACGTGGTAGACAGCCACGCAGATACACTTAAATT<br>AAaAGTAAATTTAAAAGAATACAGAAATCACATcGGAAATTTTCCTG<br>GGATGTAACTCAGCTGGCCTAGCTCCAGATCCTGGGTTCAGATGCA<br>GCGCTGCAGACAAAGAAGAAAGAAGAAAGCATTAGCCCTAGAATT<br>GCCTCCATCTTCTTCTAAATCCCTGTCCCCTCACCTACCTCCTCCTCC<br>CCATGT</td></tr>
<tr><td></td><td>+/49C</td><td>CTCAGAATAGAGAGATGGAGGCATTCTAGGGCTAATGCTTTCTTCTT<br>TCTTCTTTGTCTGCAGCGCTGCATCTGAACCCAGGATCTGGAGCTAG<br>GCCAGCTGAGTTACATCCCAGGAAAATTTCCgATGTGATTTCTGTAT<br>TCTTTTAAATTTACTTTTAATTTAAGtGTATCTGCGTGGCTGTCTACC<br>ACGTGTGtGCGGGTGCCCAGGACGGCCAGGAGAGGCTGTTGAATTC<br>CCTGGAGTTGGATTTATGGGAGGTTTTGA|GCCTGGGAACCAAACTG<br>GGGTCCTATGGAAGAGACTCTGCTGACCCAGCTCAACAGTGGCTCC<br>TAGGGTTCTGAGCCTCTCTGTCTCTGTCTCTGTTTCTCTCTTCCTAAT<br>TCATTTGTTGAGTGCATGCACACACAAGTCATGACACTCCTGCAGA<br>GATCAGGGGACAAGTTACAGGAGTCAGTTCTCTCTACCATTCTGGG<br>GATTGAACTTGGACTGTTTGGCTTGGTGGCAAGCATCTTCGCATGGC<br>GAACCATCCTGCCAGCCCTTAATTTTTATTCTCTTCTCCCTTTCTCTC</td></tr>
</table>

| | | |
|---|---|---|
| | | TCTGTCTTTGCTAGGGATTGAACTCAGAGCTTTGCATATGTAGGCAA<br>GCACTCTCCTGTTGAGCTGTATTCGAATCCCTAGATAATTCCAAATC<br>AATAGCTCTCTCATTAACTTCGGAGCATTGAGTATTATACATTTTTTT<br>TTCCTCTTGCGTATAACAGAGTAGTAAGAAAGTAAAAGAGCTACTT<br>GTCTGTTTCTACTCTAAAATTTCCAGACAGGACGGAGAAAACCCAA<br>AACAGATCTCAGGTCAAGGTTGTTGTCCTATGTGTGGTACCCTCTCT<br>TTCCCCTCCCCCATCCTTTGACCTTTGAGTGTCTGTGTATCAGTCCCG<br>TGTGACCCTGGAATGCTGCAGCCCCTCCGAGACTGTTTTCTCTGTAC<br>CATCTACACTGTCTGGATAGTTAGTTTCTGCTCTGAGATGTCTGGTT<br>CTACACACTAGCAGGAAGAGATGGcTTGTCTAGACAGTCTTCATGAT<br>ACTGAACATCACAGCGATGATGATGGGCGTCCTAAATGGAGGGCTA<br>GTTCTCATGTGATCACTATGCACGCCCTAAGGTCACCCAGATGCTCT<br>GGAGGTTCAGACTTACCTCAGGAACTAGCTGGACTGCTAGTACTCA<br>CGAGATCAATCCTATAAGAACCTCAATGCTGTAGTCTGACTCATGA<br>AGGAACTG |
| ΔLefty1 upstream | 1B/+ | GATGGGTTTAGCATTATCACTATTTTGGAGGTCGCCCAGCAAGGGA<br>TGGAGGGTCTCAAACAGAAAATGAGACCCAGTGAGACCATACT\|GA<br>TTGGGCCACTGACCACCCCTGGGTCCTTTACACTGGTCTCGAGCCAA<br>GAAAGGCAGCGCATCGTGTCAGAAGCTGCAGACTTCATTCCAGGGC<br>CCCCCTCTCCTGGTTTGGGGCCTGCTTTCCAATCTCAAGCCTGACCT<br>GGGTCAGACCTGGGCAGGAGCCATGGGGAAGGCAGCTGCTCTTCTG<br>CATAACAAAGGGTGGCCCTGCTGGAGAGCCAGGGAACACACCAGG<br>GACGAGTCTT |
| | 13B/+ | GATGAGTTTCAGGATTATCACTATTTTGGAGGTCGCCCAGCAAGGG<br>ATGGAGGGTCTCAAACAGAAAATGAGACCCAGTGAGACC\|TGACCA<br>CCCCTGGGTCCTTTACACTGGTCTCGAGCCAAGAAAGGCAGCGCAT<br>CGTGTCAGAAGCTGCAGACTTCATTCCAGGGCCCCCCTCTCCTGGTT<br>TGGGGCCTGCTTTCCAATCTCAAGCCTGACCTGGGTCAGACCTGGGC<br>AGGAGCCATGGGGAAGGCAGCTGCTCTTCTGCATAACAAAGGGTGG<br>CCCTGCTGGAGAGCCAGGGAACACACCAGGGACGAGTCTT |
| | 14B/+ | GATAGTTTCAGCATTATCACTATTTTGGAGGTCGCCCAGCAAGGGAT<br>GGAGGGTCTCAAACAGAAAATGAGACCCAGTGAGACC\|TGACCACC<br>CCTGGGTCCTTTACACTGGTCTCGAGCCAAGAAAGGCAGCGCATCG<br>TGTCAGAAGCTGCAGACTTCATTCCAGGGCCCCCCTCTCCTGGTTTG<br>GGGCCTGCTTTCCAATCTCAAGCCTGACCTGGGTCAGACCTGGGCA<br>GGAGCCATGGGGAAGGCAGCTGCTCTTCTGCATAACAAAGGGTGGC<br>CCTGCTGGAGAGCCAGGGAACACACCAGGGACGAGTCTA |
| | 24B/+ | TATGGGGTTTCAGGATTATCACTATTTTGGAGGTCGCCCAGCAAGG<br>GATGGAGGGTCTCAAACAGAAAATGAGACCCAGTGAGACCAT\|TGG<br>GCCACTGACCACCCCTGGGTCCTTTACACTGGTCTCGAGCCAAGAA<br>AGGCAGCGCATCGTGTCAGAAGCTGCAGACTTCATTCCAGGGCCCC<br>CCTCTCCTGGTTTGGGGCCTGCTTTCCAATCTCAAGCCTGACCTGGG<br>TCAGACCTGGGCAGGAGCCATGGGGAAGGCAGCTGCTCTTCTGCAT<br>AACAAAGGGTGGCCCTGCTGGAGAGCCAGGGAACACACCAGGGAC<br>GAGTCTA |
| | 53B/+ | GGCCGTGAAGGTGTCTGGATATCACTATTTTGGAGGTCGCCCAGCA<br>AGGGATGGAGGGTCTCAAACAGAAAATGAGACCCAGTGAGACCAC<br>TGAGGGAG\|GGGCCACTGACCACCCCTGGGTCCTTTACACTGGTCTC<br>GAGCCAAGAAAGGCAGCCCATCCTGTCAGAAGCTGCAGACTTCCTT<br>CCAGGGCCCCCCTCTCCTGGTTTGGGGCCTGCTTTCCATCTCACGCC<br>TGACCTGGGTCACACCTGGGCAGGAGCCATGGCGAAGGCCGCCGCT<br>CTTCTGATAACAAAGGGTGGCCCTGCTGGAGAGCCAGGGAACACAT<br>AAAGAACTATTCTTGGCC |
| | 86B/+ | TATGAGTGTTCAGGATTATCACTATTTTGGAGGTCGCCCAGCAAGG<br>GATGGAGGGTCTCAAACAGAAAATGAGACCCAGTGA\|GATTGGGCC<br>ACTGACCACCCCTGGGTCCTTTACACTGGTCTCGAGCCAAGAAAGG |

| | | |
|---|---|---|
| | | CAGCGCATCGTGTCAGAAGCTGCAGACTTCATTCCAGGGCCCCCCT<br>CTCCTGGTTTGGGGCCTGCTTTCCAATCTCAAGCCTGACCTGGGTCA<br>GACCTGGGCAGGAGCCATGGGGAAGGCAGCTGCTCTTCTGCATAAC<br>AAAGGGTGGCCCTGCTGGAGAGCCAGGGAACACACCAGGGACGAG<br>TCTACA |
| | 87B/+ | GAGGAGTGTCAGATTATCACTATTTTGGAGGTCGCCCAGCAAGGGA<br>TGGAGGGTCTCAAACAGAAAATGAGACCCAGTGAGACCA\|CTGACC<br>ACCCCTGGGTCCTTTACACTGGTCTCGAGCCAAGAAAGGCAGCGCA<br>TCGTGTCAGAAGCTGCAGACTTCATTCCAGGGCCCCCCTCTCCTGGT<br>TTGGGGCCTGCTTTCCAATCTCAAGCCTGACCTGGGTCAGACCTGGG<br>CAGGAGCCATGGGGAAGGCAGCTGCTCTTCTGCATAACAAAGGGTG<br>GCCCTGCTGGAGAGCCAGGGAACACACCAGGGACGAGTCTACAAC<br>TCGAGACCAGTGTAGAGGACCCAGGGGTGGTCGGTTGTCTCACTGG<br>CTCCCATTTTCTGTTTGAGACCCGCCATCGCGTACTGGCGACCAGCG<br>GATAGGGGGCGGCGGTGGA |
| ΔLefty1 to Lefty2 enh | +/H2 | TACGTCGGGGGTCTAGGGGGGGGGGGGGGGANNGGTAGAAGGGCA<br>GTACACTGTCTGGATAGTTAGTTTCTGCTCTGAGATGTCTGGTTCTA<br>CACACTAGCAGGGAAAGAGGATGGcTTGTCTAGACCAGTCCTTCATT<br>GATACTGAACATCACCAGGCGATGATGATG\|GGCCACTGACCACCCC<br>TGGGTCCTTTACACTGGTCTCGAGCCAAGAAAGGCAGCGCATCGTG<br>TCAGAAGCTGCAGACTTCATTCCAGGGCCCCCCTCTCCTGGTTTGGG<br>GCCTGCTTTCCAATCTCAAGCCTGACCTGGGTCAGACCTGGGCAGG<br>AGCCATGGGGAAGGCAGCTGCTCTTCTGCATAACAAAGGGTGGCCC<br>TGCTGGAGAGCCAGGGAACACACCAGGGACGAGTCTTACAATCCAC<br>TATGGTTTCTTCATCTATCCCCCGACTAAGATTGTCTGAATCTGCAA<br>ATAAAACCTCTAGATACTGGTGTAATGCAAATCCCAGGTAAACAGT<br>GACTGATATATTGTGTAACCAAGTCCCAGATTCcGTACGGGTTTTAT<br>TTGGCAGCCCTACCCCTGGCAGCTTGCTCAATTTTATTTCTCCAGCG<br>CACAAAGCCACAAATTCACACTCGGaTATCCCAGCTGTGTCTTGGAA<br>GAAGGCA |
| ΔPycr2 | C6/+ | AAATCCCTGGTCCCTCTTACTAGCTTGTATACTCTTCTCATTTACTTT<br>TCTTTTAATTCCATTTTTATATGCTGTGTGTGTTGGGGGAGGgAGATG<br>GACCTCCATGTCATAAATGCATGTGGTGTCTATTCTTCCGCCCCCCA<br>TCCCACCGCTGCCTTATGTGAGCTCCAGGGgCCAACGCTCAAGACAT<br>CAAGCTTGCTGGTGCTGCTGACTGGGCCCAGGGCCTTAAGCACACT<br>AGACACACTCTCCCACCCAGTCACTTTCCCAGCCCCCACCACTTCGA<br>TATTCCACCTGCCTTCCTAGTGATAGCAGAGCACAGGAAAGGCCAC<br>AGAGCCAGGCCCTTGGGCACCCATAACTCTCCAGTCTCTCTGGGGT<br>GTTACTTAGCAACAGTCACATACGGGATGGAAGGAGTGTGtTTATAT<br>TTCTGCACGCcATAAA\|AGTGgGTGGGGTTGACCAGAGGAAGACAAG<br>AGGTAAA |
| | D7/+ | AAACCCAGGTCCCTCTTACTAGCTTGTATACTCTTCTCATTTACTTTT<br>CTTTTAATTCCATTTTTATATGCTGTGTGTGTTGGGGGAGGgAGATG<br>GACCTCCATGTCATAAATGCATGTGGTGTCTATTCTTCCGCCCCCCA<br>TCCCACCGCTGCCTTATGTGAGCTCCAGGGgCCAACGCTCAAGACAT<br>CAAGCTTGCTGGTGCTGCTGACTGGGCCCAGGGCCTTAAGC\|TGGGG<br>TTGACCAGAGAAAAGCAGAGGTAAT |
| | G5/+ | NNNNCCAGGTCCCTCTTACTAGCTTGTATACTCTTCTCATTTACTTTT<br>CTTTTAAATTCCATTTTTATATGCTGTGTGTGTTGGGGGAGGgAGAT<br>GGACCTCCATGTCATAAATGCATGTGGTGTCTATTCTTCCGCCCCCC<br>ATCCCACCGCTGCCTTATGTGAGCTCCAGGGgCCAACGCTCAAGAC<br>ATCAAGCTTGCTGGTGCTGCTGACTGGGCCCAGGGCCTTAAGCACA<br>CTAGACACACTCTCCCACCCAGTCACTTTCCCAGCCCCCACCACTTC<br>GATATTCCACCTGCCTTCCTAGTGATAGCAGAGCACAGGAAAGGCC<br>ACAGAGCCAGGCCCTTGGGCACCCATAACTCTCCAGTCTCTCTGGG<br>GTGTTACTTAGCAACAGTCACATACGGGATGGAAGGAGTGTGtTTAT |

| | | |
|---|---|---|
| | | ATTTCTGCACGC<u>cATAAAAGG\|AGT</u>GgGTGGGGTTGACCAGAGGAAG ACAANNNNNA |
| ΔLefty1 upstream + Pycr2 | 14B Lefty1 upstream + B4/B4 | TCCCCTNCTGA\|TGTTGCTAAGTAACACCCCAGAGAGACTGGCAGAG TTATGGGTGCCCAAGGGCCTGGCTCTGTGGCCTTTCCTGTGCTCTGC TATCACTAGGAAGGCAGGTGGAATATCGAAGTGGGCTATGGTGGGA GAGTGTGTCTAGTGTGCTTAAGGCCCTGGGGCCAGTCAGCAGCACC AGCAAGCTTGATGTCTAGAGCGTTGGtCCCTGGAGCTCACATAAGGC AGCGGTGGGATGGGGGGCGGAAGAATAGACACCACATGCATTTAT GACATGGAGGTCCATCTtCCTCCCCAACACACACACAGCATATAAAAT GGAATTTAAAAGAAAAGTAAATGAGAAGAGTATACCAGCTAGTAA GAGGGGAACCTAGGTGGGGGCAGGGCAGGAGCAAACCTGGTCAGG AAAGTATACCCTNTANGGAGAGGGGAACCTAGGTGGGGCCGGGGC GGGAGCAAGACCTGTCAGGAAACNG |
| | 14B Lefty1 upstream + B11/B11 | GGGCGCCAGGTCCCTCTTNTAGCTTGTATACTCTTCTCATTTACTTTT CTTTTAAATTCCATTTTTATATGCTGTTGTGTGTTGGGGGAGGgAGAT GGACCTCCATGTCATAAATGCATGTGGTGTCTATTCTTCCGCCCCCC ATCCCACCGCTGCCTTATGTGAGCTCCAGGGgCCAACGCTCAAGAC ATCAAGCTTGCTGGTGCTGCTGACTGGGCCCAGGGCCTTAAGCACA CTAGACACACTCTCCCACCCAGTCACTTTCCCAGCCCCCACCACTTC GATATTCCACCTGCCTTCCTAGTGATAG\|TGgGTGGGGTTGACCAGA GGAAGGC |
| ΔL1-31kb | B11/+ | TCCCNCTGANGGG\|CACTGACACCCCNCGGGTCCTTTANACTGGTCT CGAGCCAAGAAAGGCAGCGCATCGTGTCAGAGGCTGCAGACTTCAT TCCAGGGCCCCCCTCTCCTGGTTTGGGGCCTGCTTTCCAATCTCAAG CCTGACCTGGGTCAGACCTGGGCAGGAGCCATGGGGAAGGCAGCTG CTCTTCTGCATAACAAAGGGTGGCCCTGCTGGAGAGCCAGGGAACA CACCAGGGACGAGTCTTACAATCCACTATGGTTTCTTCATCTATCCC CCGACTAAGATTGTCTGAATCTGCAAATAAAACCTCTAGATACTGG TGTAATGCAAATCCCAGGTAAACAGTGACTGATATATTGTGTAACC AAGTCCCAGATTCaGTACGGGTTTTATTTGGCAGCCCTACCCCTGGC AGCTTGCTCAATTTTATTTCTCCAGCGCACAAAGCCACAAATTCACA CTCGGgTATCCCAGCTGTGTTG |
| | C6/+ | AATGGGNTGAAA\|CCCTGGGGTCCTTTTAACTGGGTCTCTAAGCCAA GTAAAAGGCAGCGCATCGTGTCAGAAGCTGCAGACTTCATTCCAGG GCCCCCCTCTCCTGGTTTGGGGCCTGCTTTCCAATCTCAAGCCTGAC CTGGGTCAGACCTGGGCAGGAGCCATGGGGAAGGCAGCTGCTCTTC TGCATAACAAAGGGTGGCCCTGCTGGAGAGCCAGGGAACACACCA GGGACGAGTCTTACAATCCACTATGGTTTCTTCATCTATCCCCCGAC TAAGATTGTCTGAATCTGCAAATAAAACCTCTAGATACTGGTGTAAT GCAAATCCCAGGTAAACAGTGACTGATATATTGTGTAACCAAGTCC CAGATTCaGTACGGGTTTTATTTGGCAGCCCTACCCCTGGCAGCTTG CTCAATTTTATTTCTCCAGCGCACAAAGCCACAAATTCACACTCGGg TATCCCAGCTGTGTCTGGAAAGAAGGC |
| | E5/+ | \|<u>GACC</u>NCCCCTGGGTCCTTTACACTGGTCTCGAGCCAAGAAAGGCAG CGCATCGTGTCAGAAGCTGCAGACTTCATTCCAGGGCCCCCCTCTCC TGGTTTGGGGCCTGCTTTCCAATCTCAAGCCTGACCTGGGTCAGACC TGGGCAGGAGCCATGGGGAAGGCAGCTGCTCTTCTGCATAACAAAG GGTGGCCCTGCTGGAGAGCCAGGGAACACACCAGGGACGAGTCTTA CAATCCACTATGGTTTCTTCATCTATCCCCCGACTAAGATTGTCTGA ATCTGCAAATAAAACCTCTAGATACTGGTGTAATGCAAATCCCAGG TAAACAGTGACTGATATATTGTGTAACCAAGTCCCANANTCaGTACG GNTTTTTTTTGACAGGCCTTATTTTGGCNNCCTACCCCTTGTAGTTTN NCCNATTTTATTTCTCCAACNCACNNACNCNCAAATTCACACT |
| | E8/+ | TCAAAATGGGGCATGAACACCCNGGGTCCTTAAA\|CTGGTCTCGAGC CAAGANNGGCAGCGCATCGTGTCAGAGGCTGCAGACTTCATTCCAG GGCCCCCCTCTCCTGGTTTGGGGCCTGCTTTCCAATCTCAAGCCTGA |

<table>
<tr><td></td><td></td><td>CCTGGGTCAGACCTGGGCAGGAGCCATGGGGAAGGCAGCTGCTCTT<br>CTGCATAACAAAGGGTGGCCCTGCTGGAGAGCCAGGGAACACACC<br>AGGGACGAGTCTTACAATCCACTATGGTTTCTTCATCTATCCCCCGA<br>CTAAGATTGTCTGAATCTGCAAATAAAACCTCTAGATACTGGTGTA<br>ATGCAAATCCCAGGTAAACAGTGACTGATATATTGTGTAACCAAGT<br>CCCAGATTCaGTACGGGTTTTATTTGGCAGCCCTACCCCTGGCAGCT<br>TGCTCAATTTTATTTCTCCAGCGCACAAAGCCACAAATTCACACTCG<br>GgTATCCCAGCTGTGTGGGAAAAAAAGGGAAAANNTTGGGGATCCC<br>CAGTGTGGATTTGTGGGTTTTTGGCCTGGAGAAATAAAATTGAGCA<br>AGCTGCCAGGGGTAGGGCTGCCAAATAAAACCCGTACTGAATCTGG<br>GACTTGGTTACACAATATATCAGTCCTGTTTACCTGGGATTTGCATT<br>ACCCAGTATCTAGAGGTTTTATTTGCGATTCGACATCTTAGTCGGGG<br>GATAGATGAAGAAACATAGTGGATTGTAAGATCTCCTGGTGTGTTC<br>CTGGCTCTCGCG</td></tr>
<tr><td>ΔL1-24.5kb</td><td>B11/+</td><td>GGGGGGCTTGGACCACCTCTTAGTAAGCCCACCTGTGCAGCCAGGC<br>TAGCCTGGGGTCCCTCCTGTGGGTGCCTGTTGACAATGGAATGGTTA<br>CCATCCACTTCTCCATGGAAGGACTCTCCCATGACCTGGGAGCACCT<br>TGAGCATGCTCAGTATGTTCTTTATGGTAGAGTAACGCACAGCCTTT<br>AGTCTGACCTTGAACCTGCTAATTGAGTTTTGACATCTTTTAATCTA<br>TGAGAGTTTTGTTTGATGAGGGTTTTGTTTCTTGCTATACTGACCCA<br>AGATAGCTTTGAACTCTCATGAGAGTGGC|CCACTGACCACCCCTGG<br>GTCCTTTACACTGGTCTCGAGCCAAGAAAGGCAGCGCATCGTGTCA<br>GAAGCTGCAGACTTCATTCCAGGGCCCCCCTCTCCTGGTTTGGGGCC<br>TGCTTTCCAATCTCAAGCCTGACCTGGGTCAGACCTGGGCAGGAGC<br>CATGGGGAAGGCAGCTGCTCTTCTGCATAACAAAGGGTGGCCCTGC<br>TGGAGAGCCAGGGAACACACCAGGGACGAGTCTTACAATCCACTAT<br>GGTTTCTTCATCTATCCCCCGACTAAGATTGTCTGAATCTGCAAATA<br>AAACCTCTAGATACTGGTGTAATGCAAATCCCAGGTAAACAGTGAC<br>TGATATATTGTGTAACCAAGTCCCAGATTCaGTACGGGTTATTTTGT<br>GGGAAGAAAA</td></tr>
<tr><td></td><td>C7/+</td><td>GGGGGGCTTGGATCCACCTCTTAGTAAGCCACCTGTGCAGCCAGGC<br>TAGCCTGGGGTCCCTCCTGTGGGTGCCTGTTGACAATGGAATGGTTA<br>CCATCCACTTCTCCATGGAAGGACTCTCCCATGACCTGGGAGCACCT<br>TGAGCATGCTCAGTATGTTCTTTATGGTAGAGTAACGCACAGCCTTT<br>AGTCTGACCTTGAACCTGCTAATTGAGTTTTGACATCTTTTAATCTA<br>TGAGAGTTTTGTTTGATGAGGGTTTTGTTTCTTGCTATACTGACCCA<br>AGATAGCTTTGAACTCTCATAA|GATTGGGCCACTGACCACCCCTGG<br>GTCCTTTACACTGGTCTCGAGCCAAGAAAGGCAGCGCATCGTGTCA<br>GAAGCTGCAGACTTCATTCCAGGGCCCCCCTCTCCTGGTTTGGGGCC<br>TGCTTTCCAATCTCAAGCCTGACCTGGGTCAGACCTGGGCAGGAGC<br>CATGGGGAAGGCAGCTGCTCTTCTGCATAACAAAGGGTGGCCCTGC<br>TGGAGAGCCAGGGAACACACCAGGGACGAGTCTTACAATCCACTAT<br>GGTTTCTTCATCTATCCCCCGACTAAGATTGTCTGAATCTGCAAATA<br>AAACCTCTAGATACTGGTGTAATGCAAATCCCAGGTAAACAGTGAC<br>TGATATATTGTGTAACCAAGTCCCAGATTCaGTACGGGTTATTTTNT<br>GGGCAACAAA</td></tr>
<tr><td></td><td>+/C11</td><td>GGGGGGCTNGGACCACCTCTTAGTAAGCCCACCTGTGCAGCCAGGC<br>TAGCCTGGGGTCCCTCCTGTGGGTGCCTGTTGACAATGGAATGGTTA<br>CCATCCACTTCTCCATGGAAGGACTCTCCCATGACCTGGGAGCACCT<br>TGAGCATGCTCAGTATGTTCTTTATGGTAGAGTAACGCACAGCCTTT<br>AGTCTGACCTTGAACCTGCTAATTGAGTTTTGACATCTTTTAATCTA<br>TGAGAGTTTTGTTTGATGAGGGTTTTGTTTCTTGCTATACTGACCCA<br>AGATAGCTTTGAACTCTCATA|ATTGGGCCACTGACCACCCCTGGGT<br>CCTTTACACTGGTCTCGAGCCAAGAAAGGCAGCGCATCGTGTCAGA<br>AGCTGCAGACTTCATTCCAGGGCCCCCCTCTCCTGGTTTGGGGCCTG<br>CTTTCCAATCTCAAGCCTGACCTGGGTCAGACCTGGGCAGGAGCCA</td></tr>
</table>

| | | |
|---|---|---|
| | | TGGGGAAGGCAGCTGCTCTTCTGCATAACAAAGGGTGGCCCTGCTG<br>GAGAGCCAGGGAACACACCAGGGACGAGTCTTACAATCCACTATGG<br>TTTCTTCATCTATCCCCCGACTAAGATTGTCTGAATCTGCAAATAAA<br>ACCTCTAGATACTGGTGTAATGCAAATCCCAGGTAAACAGTGACTG<br>ATATATTGTGTAACCAAGTCCCAGATTCcGTACGGGTTATTTTTTGG<br>GAAAAAAA |
| | +/F1 | GGGGGGCTTGGATCCACCTCTTAGTAAGCCCACCTGTGCAGCCAGG<br>CTAGCCTGGGGTCCCTCCTGTGGGTGCCTGTTGACAATGGAATGGTT<br>ACCATCCACTTCTCCATGGAAGGACTCTCCCATGACCTGGGAGCAC<br>CTTGAGCATGCTCAGTATGTTCTTTATGGTAGAGTAACGCACAGCCT<br>TTAGTCTGACCTTGAACCTGCTAATTGAGTTTTGACATCTTTTAATCT<br>ATGAGAGTTTTGTTTGATGAGGGTTTTGTTTCTTGCTATACTGACCC<br>AAGATAGCTTTGAACTCTCATAAGT\|GATTGGGCCACTGACCACCCC<br>TGGGTCCTTTACACTGGTCTCGAGCCAAGAAAGGCAGCGCATCGTG<br>TCAGAAGCTGCAGACTTCATTCCAGGGCCCCCTCTCCTGGTTTGGG<br>GCCTGCTTTCCAATCTCAAGCCTGACCTGGGTCAGACCTGGGCAGG<br>AGCCATGGGGAAGGCAGCTGCTCTTCTGCATAACAAAGGGTGGCCC<br>TGCTGGAGAGCCAGGGAACACACCAGGGACGAGTCTTACAATCCAC<br>TATGGTTTCTTCATCTATCCCCCGACTAAGATTGTCTGAATCTGCAA<br>ATAAAACCTCTAGATACTGGTGTAATGCAAATCCCAGGTAAACAGT<br>GACTGATATATTGTGTAACCAAGTCCCAGATTCcGTACGGGTTATTT<br>TTTGGGGAACAAA |
| | +/F11 | GGGGGGCNTGGATCCACCTCTTAGAAGCCCACCTGTGCAGCCAGGC<br>TAGCCTGGGGTCCCTCCTGTGGGTGCCTGTTGACAATGGAATGGTTA<br>CCATCCACTTCTCCATGGAAGGACTCTCCCATGACCTGGGAGCACCT<br>TGAGCATGCTCAGTATGTTCTTTATGGTAGAGTAACGCACAGCCTTT<br>AGTCTGACCTTGAACCTGCTAATTGAGTTTTGACATCTTTTAATCTA<br>TGAGAGTTTTGTTTGATGAGGGTTTTGTTTCTTGCTATACTGACCCA<br>AGATAGCTTTGAACTC\|CAAGTTTGAGGGCAGCCTT\|GATTGGGCCAC<br>TGACCACCCCTGGGTCCTTTACACTGGTCTCGAGCCAAGAAAGGCA<br>GCGCATCGTGTCAGAAGCTGCAGACTTCATTCCAGGGCCCCCTCTC<br>CTGGTTTGGGGCCTGCTTTCCAATCTCAAGCCTGACCTGGGTCAGAC<br>CTGGGCAGGAGCCATGGGGAAGGCAGCTGCTCTTCTGCATAACAAA<br>GGGTGGCCCTGCTGGAGAGCCAGGGAACACACCAGGGACGAGTCTT<br>ACAATCCACTATGGTTTCTTCATCTATCCCCCGACTAAGATTGTCTG<br>AATCTGCAAATAAAACCTCTAGATACTGGTGTAATGCAAATCCCAG<br>GTAAACAGTGACTGATATATTGTGTAACCAAGTCCCAGATTCcGTAC<br>GGGTTATTTTGTGGGANNNNNAC |
| ΔL1-14.5kb | A8/+ | GATCCGGGANAACTAACTACAACTACACCCATTAGAACACGGTAGA<br>GTACATCTTGACAAGTAATATTAACAGCAGCGGTGGTGGCGaCGGG<br>TGGACGGATGGCAGACCTGTAAGCCCAGGGCCGAGCCTTTTGTCTG<br>CCAAGGGGATAATTTGCTCTTTGCTTTTATACTTG\|GATTGGGCCACT<br>GACCACCCCTGGGTCCTTTACACTGGTCTCGAGCCAAGAAAGGCAG<br>CGCATCGTGTCAGAAGCTGCAGACTTCATTCCAGGGCCCCCTCTCC<br>TGGTTTGGGGCCTGCTTTCCAATCTCAAGCCTGACCTGGGTCAGACC<br>TGGGCAGGAGCCATGGGGAAGGCAGCTGCTCTTCTGCATAACAAAG<br>GGTGGCCCTGCTGGAGAGCCAGGGAACACACCAGGGACGAGTCTTA<br>CAATCCACTATGGTTTCTTCATCTATCCCCCGACTAAGATTGTCTGA<br>ATCTGCAAATAAAACCTCTAGATACTGGTGTAATGCAAATCCCAGG<br>TAAACAGTGACTGATATATTGTGTAACCAAGTCCCAGATTCaGTACG<br>GGTTTTATTTGGCAGCCCTACCCCTGGCAGCTTGCTCAATTTTATTTC<br>TCCAGCGCACAAAGCCACAAATTCACACTCGGgTATCCCAGCTGTGT<br>GGAGAAGAGAGGGAAATCGGCTTGGAACCCTGGAGAAAGAATTTG<br>TGGCTGTGTGCGCTGGAAAAATAGAATTGAGCAAGCTGCAGGAGTA<br>GNCTGNANATAACACCGTACTGAATCTGGGACTTGGTTACCAATAT<br>ATCACTCCTGTTTACTGGGATTTGCATTACACAGTATCTAGAGGTTT |

| | | |
|---|---|---|
| | | TATGTGCAGATTNGACAATCTTAGTCGGGGGATAGATGAAGAAACA<br>TAGTGCATTGTAAGACTCCCTGGTGTGTTCTGGTNTCGCAGGCCCTT<br>GTTATGCAGAAAAACACTGCTTCCCATGGGTCCGNCNCGCATTGGA<br>AACNCCCCCGTGGGGGTGTGGGGGCNGG |
| | C9/+ | CCCCGGGNCAACTAATACAACNTNACCCATNGAACACGGAGAAGA<br>CATCTTGGACAAGTAATATTAACAGCAGCGGTGGTGGCGaCGGGTG<br>GACGGATGGCAGACCTGTAAGCCCAGGGCCGAGCCTTTTGTCTGCC<br>AAGGGGATAATTTGCTCTTTGCTTTTATAC\|TTGGGCCACTGACCACC<br>CCTGGGTCCTTTACACTGGTCTCGAGCCAAGAAAGGCAGCGCATCG<br>TGTCAGAAGCTGCAGACTTCATTCCAGGGCCCCCCTCTCCTGGTTTG<br>GGGCCTGCTTTCCAATCTCAAGCCTGACCTGGGTCAGACCTGGGCA<br>GGAGCCATGGGGAAGGCAGCTGCTCTTCTGCATAACAAAGGGTGGC<br>CCTGCTGGAGAGCCAGGGAACACACCAGGGACGAGTCTTACAATCC<br>ACTATGGTTTCTTCATCTATCCCCCGACTAAGATTGTCTGAATCTGC<br>AAATAAAACCTCTAGATACTGGTGTAATGCAAATCCCAGGTAAACA<br>GTGACTGATATATTGTGTAACCAAGTCCCAGATTCaGTACGGGTTTT<br>ATTTGGCAGCCCTACCCCTGGCAGCTTGCTCAATTTTATTTCTCCAG<br>CGCACAAAGCCACAAATTCACACTCGGgTATCCCAGCTGTGTTGGA<br>AGAGAAGGGAAATTGGGGGGGNATACCCCAGGGGGAAATAAGTGG<br>CTATGTGCGCTGGAGAAATAAAATTGAGCAAGCTGCCGGGGGTAGG<br>GCTGCCAATAAAACCGTACTGAATCTGGGACTTGGTTACCAATATA<br>TCACTCCTGTTTACTGGGATTTGCATTACACAGTATCTAGAGGTTTT<br>ATGTGCAGATTCGAAAATCTTAGTCGGGGGATAGATGAAGAAACCA<br>TAGTGGATTGTAAGACTCGTCTGGTGTGTTTCTGGCTTCNCAGGCCT<br>TTTGTANTCAAAAAAGAACTGCTTCCCCNGGGTCCGCGCGCCNTGG<br>GATAGACCCCCCCNGGGGTNGTGGAAGGAAGCCCCCCCCCNNAGA<br>GAGCGCGGGTNNAATNAGACTTTCN |
| | F12/+ | TACCCGGGANAACTACTTACAACTTACACCCATNGAACACGGAGTA<br>GACATTTGACAAGAATATTAACAGCAGCGGTGGTGGCGaCGGGTGG<br>ACGGATGGCAGACCTGTAAGCCCAGGGCCGAGCCTTTTGTCTGCCA<br>AGGGGATAATTTGCTCTTTGCTTTTATACTTGGATCG\|TGGGCCACTG<br>ACCACCCCTGGGTCCTTTACACTGGTCTCGAGCCAAGAAAGGCAGC<br>GCATCGTGTCAGAAGCTGCAGACTTCATTCCAGGGCCCCCCTCTCCT<br>GGTTTGGGGCCTGCTTTCCAATCTCAAGCCTGACCTGGGTCAGACCT<br>GGGCAGGAGCCATGGGGAAGGCAGCTGCTCTTCTGCATAACAAAGG<br>GTGGCCCTGCTGGAGAGCCAGGGAACACACCAGGGACGAGTCTTAC<br>AATCCACTATGGTTTCTTCATCTATCCCCCGACTAAGATTGTCTGAA<br>TCTGCAAATAAAACCTCTAGATACTGGTGTAATGCAAATCCCAGGT<br>AAACAGTGACTGATATATTGTGTAACCAAGTCCCAGATTCaGTACGG<br>GTTTTATTTGGCAGCCCTACCCCTGGCAGCTTGCTCAATTTTATTTCT<br>CCAGCGCACAAAGCCACAAATTCACACTCGGgTATCCCAGCTGTGT<br>GGGAAGAAGAGGGAAATCAGNTTGGGAGACCCGAGGGAGAATTTG<br>TGGCTTTGTGCGCTGGATAAATAAGATTGAGCAAGCTGCCGGGGTA<br>GGGCTGCANATAACACCGTACTGAATCTGGGACTTGGTTACCAATA<br>TATCACTCCTGTTTACCTGGGATTTGCATTACACAGTATCTAGAGGT<br>TTTATTTGCAGATTCGAACATCTTAGTCGGGGGATAGATGAAGAAA<br>CCATAGTGGGATTGTAAGACTCGTCTGGTGTGTTTCTGGCTTCGNGG<br>GCCCTTGTTATGCAAAAAAACACTGCTTCCCCTGGGTCCGCCCCCGC<br>GGGGGGGTCCCCCCCNCCGGGGGTGTGGGGTNAGAGCCCCCCCCCC<br>CGGNGGGGGGGGGTTTAAATA |
| | G12/+ | TTCCGGGAACAACACATACAACTACACCCATAAGAAGCACGGTAGA<br>GTACATCTCTGACAAGAAATTAACAGCAGCGGTGGTGGCGaCGGGT<br>GGACGGATGGCAGACCTGTAAGCCCAGGGCCGAGCCTTTTGTCTGC<br>CAAGGGGATA\|GCCACTGACCACCCCTGGGTCCTTTACACTGGTCTC<br>GAGCCAAGAAAGGCAGCGCATCGTGTCAGAAGCTGCAGACTTCATT<br>CCAGGGCCCCCCTCTCCTGGTTTGGGGCCTGCTTTCCAATCTCAAGC |

<table>
<tr><td></td><td></td><td>CTGACCTGGGTCAGACCTGGGCAGGAGCCATGGGGAAGGCAGCTGC<br>TCTTCTGCATAACAAAGGGTGGCCCTGCTGGAGAGCCAGGGAACAC<br>ACCAGGGACGAGTCTTACAATCCACTATGGTTTCTTCATCTATCCCC<br>CGACTAAGATTGTCTGAATCTGCAAATAAAACCTCTAGATACTGGT<br>GTAATGCAAATCCCAGGTAAACAGTGACTGATATATTGTGTAACCA<br>AGTCCCAGATTCaGTACGGGTTTTATTTGGCAGCCCTACCCCTGGCA<br>GCTTGCTCAATTTTATTTCTCCAGCGCACAAAGCCACAAATTCACAC<br>TCGGgTATCCCAGCTGTGTTGGAAAAAAAGGGGAAATTTGTGGGTT<br>NTACCCTAGGGTAAATAAATGGCTCTGTGCGCTGGAGAAATAAAAT<br>TGAGCAAGCTGCCGGCGTAGGGCTGCAAATAATACCGTACTGAATC<br>TGGGACTTGGTTACCAATATATCATTCCTGTTTACCTGGGATTTGCA<br>TTACCCAGTATCTAGAGGTTTTATTTGCAGATTCGACATCTTAGTCG<br>GGGGATAGATGAAGAAACCATAGTGGATTGTAAGACTCCCTGGTGT<br>GTTCTGGTTTCGCANGCCCTTTGTATTCAAAAAAACACTGCTTCCCA<br>TGGTCCCGCCCGCTCCGATTTAACCCCCGGCCGGGTTGGATNAGGG<br>GGCCCCCCACACCAANGAGGGGGGCCAAAAAAAAACCTCTTTCTAA<br>AAAAT</td></tr>
<tr><td></td><td>G7/+</td><td>CACGGGGAAAAATCATATAAACACACCCTAGAGAACACGGAGAGA<br>GATCTATGACAAGAAGAGTAACAGCAGCGGTGGCGGCGaCGGGTGG<br>ACGGAGGGCAGACCCGTAAGCCCAGGGCCGAGCCTTTTGTCTGCCA<br>AGGGGATAATTTGCTCTTTGCTTTTATACTTGGATCGT|GATTGGGCC<br>ACTGACCACCCCTGGGTCCTTTACACTGGTCTCGAGCCAAGAAAGG<br>CAGCGCATCGTGTCAGAAGCTGCAGACTTCATTCCAGGGCCCCCCT<br>CTCCTGGTTTGGGGCCTGCTTTCCAATCTCAAGCCTGACCTGGGTCA<br>GACCTGGGCAGGAGCCATGGGGAAGGCAGCTGCTCTTCTGCATAAC<br>AAAGGGTGGCCCTGCTGGAGAGCCAGGGAACACACCAGGGACGAG<br>TCTTACAATCCACTATGGTTTCTTCATCTATCCCCCGACTAAGATTGT<br>CTGAATCTGCAAATAAAACCTCTAGATACTGGTGTAATGCAAATCC<br>CAGGTAAACAGTGACTGATATATTGTGTAACCAAGTCCCAGATTCaG<br>TACGGGTTTTATTTGGCAGCCCTACCCCTGGCAGCTTGCTCAATTTT<br>ATTTCTCCAGCGCACAAAGCCACAAATTCACACTCGGgTATCCCAGC<br>TGGTTGGGAAAAAAGGGAAACCGGTTTGGGATCCCCAAGANTGAAT<br>TTGTGGCTTTGTGCGCTGGAGAAATAAAATTGAGCACGCTGCCGGG<br>GTAGGGCTGCAAATAAAACCGTACTGAATCTGGGACTTGGTTACCA<br>ATATATCACTCCTGTTTACCTGGGATTTGCATTACCCAGTATCTAGA<br>GGTTTTATATGCAGATTCNACATCTTAGTCGGGGGATAGATCAAGA<br>AACATAGTGGATTGTAAGACTCNCTGCTGTGTTCTGGCTTCGCAGGC<br>CCTTTTTATGCAAAAAAACACTGCTTCCCNNGGTTCCGCCCCNCGNG<br>GTGG</td></tr>
<tr><td></td><td>+/C8</td><td>TTCCGGNACAACACTACAACTACACCCATNGGAACACGGAAGAGTA<br>CATCTCTGACAAGAATATTAACAGCAGCGGTGGTGGCGgCGGGTGG<br>ACGGATGGCAGACCTGTAAGCCCAGGGCCGAGCCTTTTGTCTGCCA<br>AGGGGATAATTTGCTCTTTGCTTTTATACTTGGATC|GATTGGGCCAC<br>TGACCACCCCTGGGTCCTTTACACTGGTCTCGAGCCAAGAAAGGCA<br>GCGCATCGTGTCAGAAGCTGCAGACTTCATTCCAGGGCCCCCCTCTC<br>CTGGTTTGGGGCCTGCTTTCCAATCTCAAGCCTGACCTGGGTCAGAC<br>CTGGGCAGGAGCCATGGGGAAGGCAGCTGCTCTTCTGCATAACAAA<br>GGGTGGCCCTGCTGGAGAGCCAGGGAACACACCAGGGACGAGTCTT<br>ACAATCCACTATGGTTTCTTCATCTATCCCCCGACTAAGATTGTCTG<br>AATCTGCAAATAAAACCTCTAGATACTGGTGTAATGCAAATCCCAG<br>GTAAACAGTGACTGATATATTGTGTAACCAAGTCCCAGATTCcGTAC<br>GGGTTTTATTTGGCAGCCCTACCCCTGGCAGCTTGCTCAATTTTATTT<br>CTCCAGCGCACAAAGCCACAAATTCACACTCGGaTATCCCAGCTGTG<br>TGGGAGAGAAGAGGGAAATTCGGGGGGGGNNANCCAAGNNAAAAT<br>ATGTGGCTATGTGCGCTGGAAAAATAAGATTGAGCAAGCTGCCAGG<br>GGTAGGGCTGCCAAATAAAACCGTACGGAATCTGGGACTTGGTTAC</td></tr>
</table>

<table>
<tr><td></td><td></td><td>CAATATATCACTCCTGTTTACTGGGATTTGCATTACACAGTATCTAG<br>AGGTTTTATGTGCAGATTCGAACATCTTAGTCGGGGGATAGATGAA<br>GAAACCATAGTGGATTGTAAGACTCGTCTGCTGTGTTTCTGGCTTCG<br>CGGGCCCTTTTTATGCAAAAAAGCACTGCTTCCCCTGGGTCGCCGCG<br>CCCGGGGTGGCCCCCCCCCGGTGGGTGGGGTGGGGAGGCCCCCCCC<br>CCC</td></tr>
<tr><td></td><td>+/H2</td><td>TTCCGGGANAACTAATTACAACNTAACCCATGGGAACACGGAAGTA<br>GTACATCTTGTACAAGAATATTAACAGCAGCGGTGGTGGCGgCGGG<br>TGGACGGATGGCAGACCTGTAAGCCCAGGGCCGAGCCTTTTGTCTG<br>CCAAGGGGATAATTTGCTCTTTGCTTTTATACTTGGATT|GGCCACTG<br>ACCACCCCTGGGTCCTTTACACTGGTCTCGAGCCAAGAAAGGCAGC<br>GCATCGTGTCAGAAGCTGCAGACTTCATTCCAGGGCCCCCCTCTCCT<br>GGTTTGGGGCCTGCTTTCCAATCTCAAGCCTGACCTGGGTCAGACCT<br>GGGCAGGAGCCATGGGGAAGGCAGCTGCTCTTCTGCATAACAAAGG<br>GTGGCCCTGCTGGAGAGCCAGGGAACACACCAGGGACGAGTCTTAC<br>AATCCACTATGGTTTCTTCATCTATCCCCCGACTAAGATTGTCTGAA<br>TCTGCAAATAAAACCTCTAGATACTGGTGTAATGCAAATCCCAGGT<br>AAACAGTGACTGATATATTGTGTAACCAAGTCCCAGATTCcGTACGG<br>GTTTTATTTGGCAGCCCTACCCCTGGCAGCTTGCTCAATTTTATTTCT<br>CCAGCGCACAAAGCCACAAATTCACACTCGGaTATCCCAGCTGTGC<br>GGGNGAAAAAAGGGAAATCTGGTTGGGATATCTTAGTGAGAAATTT<br>GTGGGTTTGTGCGCTGGATAAATAAGATTGAGCAAGCTGCCAGGGG<br>TAGGGCTGCCANTAAAACCGTACGGAATCTGGGACTTGGTTACCAA<br>TATATCACTCCTGTTTACTGGGATTTGCATTACACCGTATCTAGAGG<br>TTTTATGTGCAGATTCGAAAATCTTAGTCGGGGGATAGATGAAGAA<br>ACCATAGTGGATTGTAAGACTCGTCTGGTGTGTTCTGGCTTCGCAGG<br>CCCTTTGTATGCAAAAAAGCACTGCTTCCCCNGGGTTGCCTCCCGNC<br>GGGNGT</td></tr>
<tr><td></td><td>+/G4</td><td>TCCCGGNANAACTACTTAAACTTACACCCATGGAACACGGTAGTAA<br>GACATCTTGACAAGAATATTAACAGCAGCGGTGGTGGCGgCGGGTG<br>GACGGATGGCAGACCTGTAAGCCCAGGGCCGAGCCTTTTGTCTGCC<br>AAGGGGATAATTTGCTCTTTGCTTTTATACTTG|GATTGGGCCACTGA<br>CCACCCCTGGGTCCTTTACACTGGTCTCGAGCCAAGAAAGGCAGCG<br>CATCGTGTCAGAAGCTGCAGACTTCATTCCAGGGCCCCCCTCTCCTG<br>GTTTGGGGCCTGCTTTCCAATCTCAAGCCTGACCTGGGTCAGACCTG<br>GGCAGGAGCCATGGGGAAGGCAGCTGCTCTTCTGCATAACAAAGGG<br>TGGCCCTGCTGGAGAGCCAGGGAACACACCAGGGACGAGTCTTACA<br>ATCCACTATGGTTTCTTCATCTATCCCCCGACTAAGATTGTCTGAAT<br>CTGCAAATAAAACCTCTAGATACTGGTGTAATGCAAATCCCAGGTA<br>AACAGTGACTGATATATTGTGTAACCAAGTCCCAGATTCcGTACGGG<br>TTTTATTTGGCAGCCCTACCCCTGGCAGCTTGCTCAATTTTATTTCTC<br>CAGCGCACAAAGCCACAAATTCACACTCGGaTATCCCAGCTGTGTG<br>GGAAAAAAAGGGAAATCCGGCTTGGGAGAGCCCGAGNGAGAATTT<br>GTGGCTTTGTGCGCTGGAAAAATAAGATTGAGCAAGCTGCCGGAGT<br>AGGGCTGCCAATAAAACCGTACGGAATCTGGGACTTGGTTACCAAT<br>ATATCACTCCTGTTTACCTGGGATTTGCATTACACAGTATCTAGAGG<br>TTTTATTTGCAGATTCGACATCTTAGTCGGGGGATAGATGAAGAAA<br>CATAGTGGATTGTAAGACTCTCTGGTGTGTTCTGGCTTCGNGGGCCC<br>TTGTTATGCNAAAAAGCACTGCTTCCCCTGGTTTCCTCGCCCTCCGG<br>NGTTGCCCCCCCCGCGTGGGGNGGNGGNGGGNCCCCCCCCNNNG<br>GGGGGGGGGGCATAAAAAAT</td></tr>
<tr><td></td><td>+/F9</td><td>TCCCGGCACAACTAATTACAACTTACACCCATTGGAACACGGTAGT<br>AGACATTTGACAAGTAATATTAACAGCAGCGGTGGTGGCGgCGGGT<br>GGACGGATGGCAGACCTGTAAGCCCAGGGCCGAGCCTTTTGTCTGC<br>CAAGGGGATAATTTGCTCTTTGCTTTTATACTTGGATCGCT|GATTGG<br>GCCACTGACCACCCCTGGGTCCTTTACACTGGTCTCGAGCCAAGAA</td></tr>
</table>

| | | |
|---|---|---|
| | | AGGCAGCGCATCGTGTCAGAAGCTGCAGACTTCATTCCAGGGCCCC<br>CCTCTCCTGGTTTGGGGCCTGCTTTCCAATCTCAAGCCTGACCTGGG<br>TCAGACCTGGGCAGGAGCCATGGGGAAGGCAGCTGCTCTTCTGCAT<br>AACAAAGGGTGGCCCTGCTGGAGAGCCAGGGAACACACCAGGGAC<br>GAGTCTTACAATCCACTATGGTTTCTTCATCTATCCCCCGACTAAGA<br>TTGTCTGAATCTGCAAATAAAACCTCTAGATACTGGTGTAATGCAA<br>ATCCCAGGTAAACAGTGACTGATATATTGTGTAACCAAGTCCCAGA<br>TTCcGTACGGGTTTTATTTGGCAGCCCTACCCCTGGCAGCTTGCTCA<br>ATTTTATTTCTCCAGCGCACAAAGCCACAAATTCACACTCGGaTATC<br>CCAGCTGTGTGGGAAAAAAAGGGAAATCCGGCTGGGAGNCCCGAG<br>TGAGAATANGTGGCTATGTGCGCTGGATAAATAAGATTGAGCAAGC<br>TGCCAGGAGTAGGGCTGCCAAATAACACCGTACGGAATCTGGGACT<br>TGGTTACCAATATATCACTCCTGTTTACTGGGATTTGCATTACACAG<br>TATCTAGAGGTTTTATTTGCAGATTCAACATCTTAGTCGGGGGATAG<br>ATAAAGAAACCATAGTGGATTGTAAGACTCGTCTGCTGTGTTCCGG<br>CTTCGNGGGCCCTTTTTATGCAAAAAGCACTGCTTCCCCTNTGGTCC<br>TCCACGGGAGAGATNACCCACCCGGAGGGTGGTGTTGGATAAACCC<br>CCCCCCCCAGAGGGGGGGGCGTAAAGAACCCTCC |

**S2 Table: Guide RNA sequences for CRISPR/Cas9 mediated deletions.**

| Target Region | Left Guide Sequence | Right Guide Sequence |
|---|---|---|
| ΔLefty1enh | ATATCCCCCGCAAGACAGTA | CAGGAGCTCACAGGGCCACA |
| ΔLefty2enh | TTCCCAGGCTCCGCATCAGG | TGATGGGGCGTTCCCTAAAT |
| ΔLefty1 upstream | ATATCCCCCGCAAGACAGTA | GGTCAGTGGCCCAATCACGA |
| ΔLefty1 to Lefty2 enh | TGATGGGGCGTTCCCTAAAT | GGTCAGTGGCCCAATCACGA |
| ΔPycr2 | GCGAGCGGGGCAGACCAGAG | GCACGCCATAAAAGGAACAG |
| ΔL1-31kb | GCGAGCGGGGCAGACCAGAG | GGTCAGTGGCCCAATCACGA |
| ΔL1-24.5kb | CTTTGAACTCTCATAAGCCA | GGTCAGTGGCCCAATCACGA |
| ΔL1-14.5kb | CTTTTATACTTGGATCGGGT | GGTCAGTGGCCCAATCACGA |

**S3 Table: QPCR primers for gene expression analysis (SNPs indicated as lowercase)**

| mRNA | Allele | Forward Sequence | Reverse Sequence |
|---|---|---|---|
| Lefty1 | 129 | TCATGCAAATCTGAAGTGCTCa | GGACAGAGACCACTGGGTCA |
| Lefty1 | CAST | CATGCAAATCTGAAGTGCTCcT | GGACAGAGACCACTGGGTCA |
| Lefty2 | 129 | GCAAGAGGTTCAGCCAGAAt | CACACACAAAGAAAGGGTCtACAt |
| Lefty2 | CAST | GCAAGAGGTTCAGCCAGAAc | ACACAAAGAAAGGGTCcACAc |
| Pycr2 | 129 | CAGCTCAGGGAAACTCCTTACa | GTGGCACCAGGTGAACAGAT |
| Pycr2 | CAST | CAGCTCAGGGAAACTCCTTACc | GTGGCACCAGGTGAACAGAT |
| Sdha | n/a | ACTGGGATGGGCTCCTTAGT | GCCCTGAGAAAGATCACGTC |
